# Qimai: a multi-agent framework for zero-shot DNA-protein interaction prediction

**DOI:** 10.1101/2025.09.30.679628

**Authors:** Cong Liu, Mina Yao, Wei Wang

## Abstract

Accurate prediction of DNA-protein interactions, a fundamental task in genomics, is limited by the poor generalization of existing models to novel proteins not seen during training. To address this challenge, we introduce Qimai, a modular AI agent framework that integrates deep learning predictions with biological evidence using Large Language Model (LLM) as reasoning engine. Qimai combines direct motif evidence from the query protein, indirect motif evidence from its interactors, and quantitative prediction from a new transformer-based DPI model to produce explainable predictions with confidence scores. On a benchmark of 78 unseen proteins, Qimai consistently outperforms standalone deep learning models across all metrics, increasing the Area Under Curve of the Precision-Recall (AUC-PR), the Area Under Curve of the Receiver Operating Characteristic (AUC-ROC), and Matthews Correlation Coefficient (MCC) by 17.6%, 15.6%, and 244% respectively compared to the best standalone model. Ablation analyses reveal that this gain is driven by the LLM’s ability to dynamically weigh diverse evidence, with indirect motif evidence of co-factors particularly critical for unseen proteins. Qimai establishes a generalizable and interpretable paradigm for integrating heterogeneous data in predictive genomics. This framework is accessible via the Qimai web portal (https://qimai.wanglab.ucsd.edu/).

## Introduction

Transcription factors (TFs) bind to specific DNA sequences to control transcriptional programs, which are essential for cellular functions. While TF binding is partially determined by short sequence motifs, it is also highly context-dependent, modulated by local chromatin state, co-binding factors, and protein– protein interactions (PPIs)^1–3^. This complexity poses a persistent challenge for accurately predicting TF binding sites and generalizing these predictions to any protein from DNA sequence alone.

Deep learning has recently transformed this field. Models such as DeepSEA^4^, Sei^5^, and DNABERT^6^ have leveraged large-scale ChIP-seq datasets to learn complex sequence-to-binding relationships. However, these models are often limited by a lack of interpretability and an inability to generalize beyond their training data, particularly to novel TFs not seen during training^7^. A critical challenge arises because many proteins detected in ChIP-seq do not bind DNA intrinsic sequence specificity. Instead, they are recruited by other DNA-binding proteins or function within larger complexes that assemble on chromatin. These indirect but functionally indispensable associations are fundamental to transcriptional regulation but are beyond the prediction capability of the existing models.

For example, a master regulator like RUNX1 often co-localizes with its binding partners TAL1 and GATA proteins to form a regulatory circuit, controlling pluripotency and self-renewal^8^. Likewise, structural proteins such as cohesin components (e.g., RAD21) are often co-precipitated with sequence-specific factors like CTCF, forming a complex critical for higher-order genome folding, stabilizing enhancer–promoter loops, and shaping the regulatory landscape^9^. Similarly, many chromatin-modifying enzymes lack intrinsic DNA-binding ability but are guided to precise loci by sequence-specific TFs^10^. These examples highlight that predicting DNA–protein interactions requires capturing not only motif-driven binding but also the recruitment of non-sequence-specific proteins essential for 3D genome organization, chromatin state maintenance, and signal integration. Importantly, by failing to disentangle these mechanisms, models become uninterpretable and cannot generalize to novel interactions mediated by proteins, creating a critical need for a new framework that can reason about these distinct modes of interaction.

Here, we introduce Qimai (**Q**uantitative **I**nference of **M**olecular interactions by **AI** agents), a multi-agent AI system for predicting DNA–protein interactions that integrates deep learning predictions with diverse biological evidence to improve generalization and interpretability. Our method consists of four distinct yet cooperative agents: a Decision Agent, a prediction model trained on ChIP-seq data; an Evidence Retriever, which determines whether the input sequence contains motifs of the target TF or of its known protein-protein interaction (PPI) partners; and a Reasoning Agent and a Critique Agent, which together integrate predictions and motif context to infer interactions, explains the reasoning and assesses prediction confidence. This modular structure mimics human expert reasoning: a quantitative prediction from the deep learning model is reinforced by the presence of the TF’s own motif in the DNA, or, if absent, by motifs of its known interactors. This indirect evidence is especially informative for proteins like novel TFs without known motifs or cohesion and chromatin modifiers, which lack intrinsic sequence motifs but exert central regulatory roles through recruitment by DNA-binding partners.

Across a range of benchmarks, Qimai significantly outperforms state-of-the-art models, including DeepSEA^4^, Sei^5^, and DNABERT^6^—in both standard and zero-shot settings, accurately predicting binding of previously unseen proteins on unseen DNA sequences. By disentangling sequence modeling from prior biological knowledge, and recombining them in a transparent decision framework, our architecture represents a conceptual shift in regulatory modeling: from end-to-end black boxes to modular, interpretable agents informed by biological reasoning.

## Results

### The Qimai Framework: LLM-Driven Synthesis of Heterogeneous Evidence

To address the challenge of predicting DNA-protein interactions involving unseen proteins (i.e. proteins not included in the training data), we developed Qimai, a multi-agent framework that employs Large Language Model (LLM) to synthesize predictions from a deep learning model with direct and indirect motif evidence (**Figure 1A**). We used Gemini-2.0-flash for performance evaluation and also assessed other LLMs (see **Supplementary Note B**). The Decision Agent features a new dual-input transformer DPI model (referred to as DPI model hereafter) that leverages pre-trained DNABERT and AlphaFold2 embeddings, enabling generalization to unseen DNA sequences and proteins (**Figure 1B**). The Evidence Retriever Agent then collects relevant evidences by: (1) scanning the DNA sequence for direct binding motifs, if known, of the query protein; (2) identifying motifs from known protein interactors (i.e. indirect motif evidence) whose bindings suggest possible complex formation with the query protein; and (3) generating a baseline prediction from the deep learning model such as the transformer DPI model we developed in this work (**Supplementary Figure S1A**). This unified evidence is passed to an LLM Reasoning Agent, which evaluates the evidence to produce a final, explainable prediction, and a Critique Agent, which assigns a rule-based confidence score.

**Figure 1:**
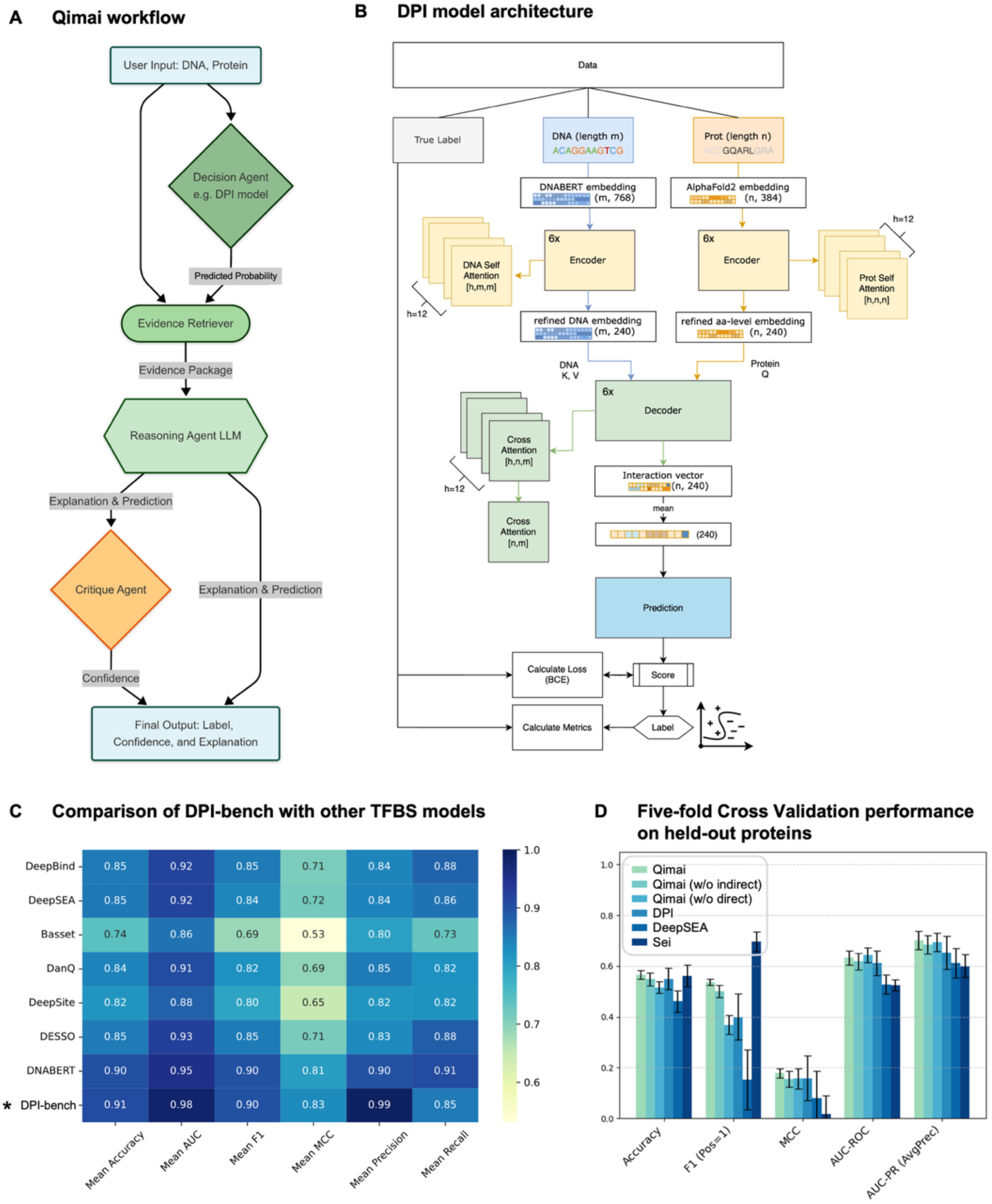
Qimai framework and superior generalization performance on unseen proteins. **(A) Overall workflow.** User input (DNA sequence, protein name) initiates the Evidence Retriever and Decision Agent. Retrieved motif evidence along with predicted probability is passed to a Reasoning Agent LLM, which generates an explanation and prediction. A Critique Agent then assigns a confidence score to produce the final output. **(B) DPI model architecture.** We developed a new transformer-based deep learning model (DPI model) as one source of evidence. DNA and protein sequences are embedded (e.g., via DNABERT and AlphaFold2-derived embeddings, respectively) and processed by separate encoders. A cross-attentional decoder then integrates these representations to predict an interaction score and label. The true label is used for training via loss calculation (e.g., Binary Cross-Entropy, BCE) and metric evaluation. h: number of attention heads; m, n: sequence lengths; embedding dimensions are shown in parentheses. **(C) Benchmark DPI model with the state-of-the-art models.** DPI model showed clear improvement over other models in almost every evaluation metrics, highlighting the effectiveness of pre-trained DNA embeddings. **(D) Five-fold Cross Validation performance within Encode3and4 dataset**. The full Qimai system (lightest green) is compared against standalone deep learning models (DeepSEA, DPI, Sei) and ablated versions of the system. “Qimai (w/o direct)” and “Qimai (w/o indirect)” represents removing direct and indirect motif information from full Qimai respectively. This annotation is used throughout the paper. Results are averaged over five-fold cross validation results with error bars showing standard deviation.

### The new transformer DPI model achieves superior performance

We first compared the new transformer DPI model with the existing ones including DeepBind, DeepSEA, and DNABERT (**Figure 1C**). We used the ChIP_690 dataset, a standard benchmark dataset on which the other models were trained. It contains 690 TF ChIP-seq experiments from the ENCODE database processed with the DeepBind^11^ protocols. Instead of training a separate model for each experiment as DeepBind, we combined all ChIP-seq experiments together and trained a unified model. DPI model showed clear improvement over other models in almost every evaluation metric (**Figure 1C**), highlighting the effectiveness of pre-trained DNA embeddings. While DPI and DNABERT showed similar performances across most metrics, DPI achieved higher precision and lower recall, indicating our model returned fewer, yet more accurate positive predictions.

Furthermore, we extracted the self-attention weights from protein module and compared the high-attention regions with the annotated DNA-binding domains (DBDs). Thirty-two out of 34 proteins that have documented DBDs in UniProt showed at least 50% overlapped regions (**Supplementary Figure 1C**). For example, DBDs of FOXA2 and PAX5 are marked with significantly high attention. RXRA, on the other hand, has two high-attention regions: 133-159 and 261-430. The first region is related to DBD (135-200) while the second overlaps with nuclear receptor ligand-binding domain (LBD) (183-417). It demonstrated that pre-trained AlphaFold embeddings have encoded protein binding information, facilitating prediction of DNA-protein interactions.

### Qimai achieves robust performance on internal cross-validation

For a rigorous assessment, we generated five random train/test splits from Encode3and4 dataset, which is a larger and more diverse dataset including 1877 ChIP-seq experiments across 844 TFs, each time holding out 44 unseen proteins (**Methods**). When evaluated on the complete set of held-out proteins (averaged across the five splits), the full Qimai system clearly outperformed all other models (**Figure 1D**). It achieved a top F1 score of 0.537 and MCC of 0.178, a performance lift of approximately 35% and 13% over the best standalone model, DPI (F1= 0.399, MCC=0.158). The other standalone models exhibited distinct and often extreme biases: DeepSEA was highly conservative (high Specificity, low Recall), while Sei was overly optimistic (low Specificity, high Recall) (**Supplementary Table 12**). Importantly, Qimai’s performance was remarkably more stable. For the F1 score, Qimai’s standard deviation was only 0.012, nearly eight times smaller than DPI’s deviation of 0.091. Similarly, its MCC deviation (0.018) was almost five times smaller than that of DPI (0.088).

To dissect the contribution of each evidence type, we evaluated ablated versions of Qimai framework. Withholding direct motif (“Qimai (w/o direct)”) resulted in the most substantial performance decrease, with the F1 score falling from 0.537 to 0.368. Similarly, exclusion of indirect motif evidence (“Qimai (w/o indirect)”) also dropped the predictive power (F1 = 0.501). This indicated that both direct and indirect motif information provide unique, synergistic benefits for achieving optimal prediction.

Within each held-out set, we further identified “novel” proteins, defined as having less than 30% sequence identity to any protein in the corresponding 800-protein training set. This resulted in five distinct sets of novel proteins (6-12 per split) for evaluation (**Supplementary Figure S4A**). On these truly novel proteins, Qimai’s advantage was even more evident (**Supplementary Figure S4B; Supplementary Table 13**). Here, Qimai achieved an F1-score of 0.497 and MCC of 0.272, while the best standalone model, the transformer DPI, only managed an F1-score of 0.343 and MCC of 0.114, representing a much larger performance lift of nearly 45% and 138%, respectively.

### Qimai Achieves Superior Generalization on Novel Proteins in external dataset

To enable a more powerful prediction model, we re-trained the transformer DPI model on the whole dataset Encode3and4. Note that this re-trained DPI model on Encode3and4 did not see ChIP_690 data at all. We next incorporated this DPI model into Qimai and evaluated the full Qimai pipeline’s performance on the external ChIP_690 dataset, which contains unseen DNA sequences paired with 80 proteins present (i.e. seen proteins) and 78 proteins not present in the training dataset (i.e. unseen proteins). Here, DNA sequence-based models like Sei achieved strong performance (**Supplementary Figure S2A**). Qimai system remained competitive in this setting, suggesting that DNA decoding dominates predictions for known proteins.

A method’s utility for novel discovery lies in its generalization ability to new proteins. We thus performed a more stringent external validation on the 78 unseen proteins in ChIP_690. Qimai’s methodological advantage was clearly demonstrated here (**Figure 2A; Supplementary Table 7**). Qimai achieved the highest performance across all key metrics, including an AUC-PR of 0.701, MCC of 0.308 and an F1-score of 0.552. This represents a substantial improvement over the best standalone model, transformer DPI, which had an AUC-PR of 0.596, MCC of 0.090 and an F1-score of 0.322. The other standalone models, DeepSEA and Sei, struggled significantly, with Sei’s performance falling to near-random levels (AUC-PR 0.498, MCC −0.260), highlighting their limited ability to generalize to new proteins. Crucially, the final confidence score produced by Qimai is not only highly discriminative, reflected by high AUC-ROC and AUC-PR, but also well-calibrated (i.e. the confidence level corresponds to the actual likelihood of a correct prediction), confirming that the score’s value is a reliable proxy for the actual probability of a correct prediction (**Supplementary Figure S2B**).

**Figure 2:**
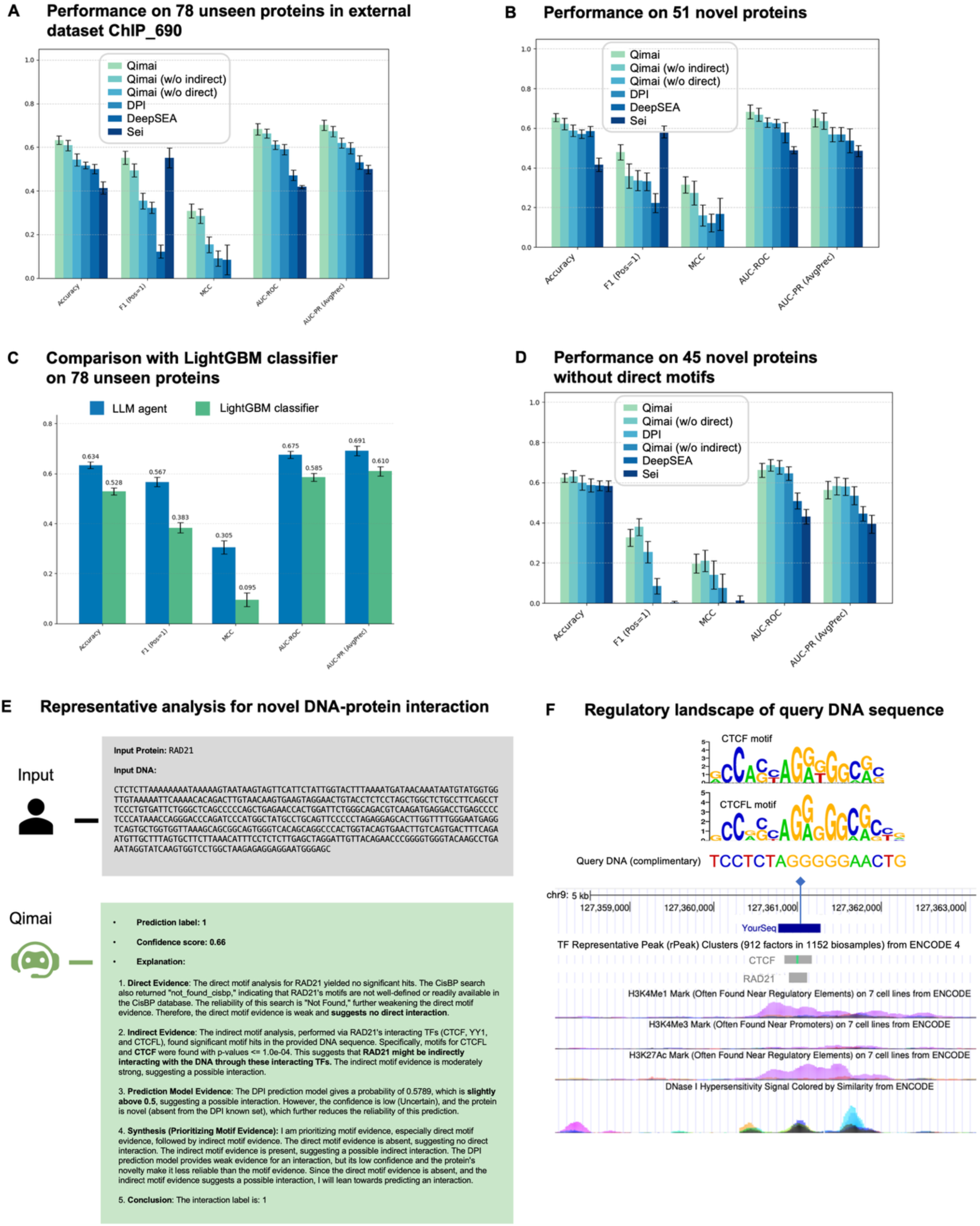
Qimai achieves superior generalization on unseen proteins by integrating motif evidence. **(A) Performance benchmark on 78 unseen proteins in ChIP_690 dataset.** The full Qimai system consistently outperforms all standalone models (DPI, DeepSEA, Sei) and ablated configurations across all metrics, with error bars representing the standard deviation over ten runs. Each run was evaluated on 500 randomly sampled pairs. **(B) Performance on 51 novel proteins.** The evaluation results from the “unseen” benchmark in Figure 2A were recalculated to include only the 51 proteins classified as novel (sequence similarity < 30%). The full Qimai system maintains the best performance, particularly in F1 and MCC. **(C) Performance of the LLM Agent versus LightGBM classifier on 78 unseen proteins.** The LLM Agent (blue) and a highly optimized LightGBM baseline (green) were evaluated on a hold-out test set of 5,000 protein-DNA pairs containing only 78 unseen proteins. Error bars represent 95% confidence intervals estimated via bootstrapping. The LLM Agent consistently outperforms the strong classifier, with the most significant relative improvement seen in the MCC. **(D) Performance on 45 novel proteins without direct motifs.** The evaluation results were recalculated from the “unseen” benchmark in Figure 2A to include only the 45 proteins classified as both novel and having no known direct motifs. Qimai’s robust performance, driven by the “Qimai (w/o direct)” configuration, is maintained on this most challenging scenario while standalone models like DeepSEA and Sei collapse to near-random levels. The failure of “Qimai (w/o indirect)” negative control confirms that indirect evidence is the primary driver of success in this *de novo* prediction scenario. **(E) Representative analysis for a novel DNA-protein interaction.** Qimai’s output for the non-canonical DNA-binding protein RAD21. Faced with no direct motif hits and an uncertain DPI model prediction, the agent leverages strong indirect evidence from RAD21’s known interactors (CTCF, CTCFL) to correctly predict an interaction with high confidence, demonstrating the framework’s evidence synthesis capabilities in a real-world scenario. **(F) Regulatory landscape of query DNA sequence.** Genomic tracks from the UCSC Genome Browser are shown. In the TF rPeaks track, the shade of the gray box is proportional to the maximum ChIP-seq signal observed across samples. The motif site is highlighted in green. The lower tracks display histone marks, where peak heights in the H3K4Me1, H3K4Me3, and H3K27Ac tracks correspond to the level of histone mark enrichment. The DNase track shows DNaseI hypersensitivity signal, indicating the open chromatin regions.

This superior performance was confirmed when focusing specifically on the novel proteins, defined as having less than 30% sequence identity to any protein in the training set of 844 Encode3and4 proteins. In total, 51 novel proteins out of 78 unseen proteins (65%) are identified (**Supplementary Figure S2C**), which are diverse across multiple TF families. Almost half of the novel proteins are also in novel protein families (23/51) (**Supplementary Figure S2D**). In this test, the full Qimai system consistently achieved highest scores across most of the key metrics (**Figure 2B**). This represents a substantial improvement over the standalone deep learning methods. For instance, Qimai’s MCC (0.314) is more than double that of the standalone model DPI’s (0.122), DeepSEA (0.166), while Sei’s negative MCC (−0.176) indicates its predictions were worse than random guessing. The standalone models exhibited distinct and often extreme biases: DeepSEA was highly conservative (high Specificity, low Recall), while Sei was overly optimistic (low Specificity, high Recall) (**Supplementary Table 8**). This demonstrates that Qimai’s ability to integrate biological motif evidence is critical for truly novel protein prediction.

Ablation studies revealed that this gain of performance stems from Qimai’s ability to integrate diverse evidence. While removing direct motif evidence (“Qimai (w/o direct)”) causes the most significant drop in performance, demonstrating its high impact when available, the removal of indirect motif evidence (“Qimai (w/o indirect)”) also consistently degraded performance compared to the full system. This confirms that both evidence are essential and complementary for optimal prediction. The consistent positive contribution of indirect evidence is particularly noteworthy, as it provides a robust and alternative source of biological information. These results underscore Qimai’s ability to intelligently integrate biologically grounded motif information via LLM-based reasoning, offering critical resilience when computational models operate outside their training domain.

### LLM Agent Outperforms a state-of-the-art Classifier through Context-aware Reasoning

We next investigated whether LLM outperformed ensemble classifier for integrating information. We compared the LLM Agent against a highly optimized LightGBM ensemble classifier on a challenging hold-out test set of 5,000 protein-DNA interactions composed of 78 unseen proteins and unseen DNAs in the ChIP_690 dataset. LightGBM performed well, achieving an AUC-ROC of 0.817 on the validation set. As any classifier, the feature weights in LightGBM are fixed and the most important contribution comes from the transformer DPI model.

Compared to LightGBM, the LLM Agent demonstrated consistently superior performance across all evaluation metrics (**Figure 2C**). The most substantial improvement was observed in MCC, where the LLM Agent achieved a score of 0.305, representing a 221% relative improvement over the baseline’s score of 0.095. This indicates a significantly better classification balance and highlights a key limitation of standard classification models, whose feature weights are fixed and cannot adjust evidence weights based on different contexts, such as weighing indirect evidence more if the deep learning model has a borderline prediction or a novel protein does not have any known motif or bind to DNA directly. Indeed, LightGBM applied the same weights learned from the training data to the unseen or novel proteins, which led to poor performance.

The LLM Agent’s performance stemming from its flexible, context-aware reasoning is exemplified in a case study of AP2A protein (**Supplementary Figure S2E**). Here, the agent was faced with strong motif evidence that contradicted a low-probability deep learning model prediction. The LightGBM classifier, misled by its learned dependence on the model score, failed. In contrast, the LLM Agent correctly identified the conflict, evaluated the quality of the evidence, and prioritized the strong motif evidence to reach the correct conclusion. Additional examples are shown in **Supplementary Note D**.

To further validate the inferential process, we conducted an “LLM-as-a-Judge” evaluation where three independent LLM judges (phi4, deepseek-r1, and qwen3) assessed 500 prediction explanations generated by Qimai. This cross-examination confirmed consistently high quality of the agent’s reasoning across all evaluation criteria (**Supplementary Figure S2F**). All three judges awarded near-perfect scores for key capabilities such as Factual Accuracy (avg. 4.99/5), Conflict Resolution (avg. 4.97/5), and Overall Quality (avg. 4.95/5), indicating that the generated explanations are robust, coherent, and logically sound. Agreement between diverse judges confirms the objective quality of the outputs and indicates that the Qimai’s superior performance is a direct result of its robust, evidence-based inferential capabilities. This capacity to dynamically weigh and synthesize conflicting data streams, much like a human expert, marks a significant advantage of the agent-based framework.

### Indirect Evidence is Critical for Predicting de novo Interactions

The importance of indirect evidence is most apparent when evaluating its performance on the most challenging subset of proteins: 45 novel proteins (sequence similarity < 30%) that also lack known direct binding motifs. This scenario represents a true *de novo* discovery problem where methods relying on direct motif information or protein-specific training data are expected to fail. It is not surprising that the standalone Sei and DeepSEA models collapsed to near-random performance on this cohort (**Figure 2D; Supplementary Table 9**).

In stark contrast, Qimai demonstrated remarkably robust predictive power, a success driven almost entirely by indirect evidence. The configuration relying only on indirect motif evidence (“Qimai (w/o direct)”) achieves the highest performance, with a MCC of 0.211, nearly triple that of the negative control (“Qimai (w/o indirect)”, MCC = 0.076). The standalone transformer DPI model, while also outperforming Sei and DeepSEA due to its better generalization, still falls significantly short of the Qimai configurations that include indirect evidence. This observation decisively proves that Qimai’s ability to leverage the indirect evidence pathway is the primary driver of its success in these challenging cases. By identifying interacting TFs and their motifs, the LLM agent can successfully reason about potential interactions even when all direct information about the query protein is absent, a critical capability for expanding our understanding of uncharacterized proteins.

Indirect motif evidence proves especially critical for predicting interactions involving non-canonical transcription factors (non-TFs) or TFs without known direct motifs. In these cases, indirect evidence by scanning for motifs of the query protein’s PPI partners provides crucial, otherwise missing, clues about potential DNA binding mediated through protein complex or co-factor recruitment. To quantify the impact of this evidence, we analyzed 4713 correct predictions involving 186 unique non-TF proteins or TFs without known motifs: 76.8% were “rescued” or strongly supported by significant indirect motif hits, demonstrating its decisive, not merely supplementary, role, especially for proteins with limited or no direct binding motif information.

A representative example is RAD21, a cohesin complex component rather than a TF, that does not have a known motif in CisBP (**Figure 2E**). For example, for a query DNA sequence at chr9:127,360,766-127,361,277, which is characterized by strong epigenetic signals indicative of an active regulatory element, including enrichment for the active enhancer histone marks H3K27ac and H3K4me1, as well as high DNase I hypersensitivity, signifying open chromatin (**Figure 2F**). The DPI model only produced a borderline prediction (0.5789). However, Qimai’s Evidence Retriever identified strong motif matches for RAD21’s key interactors, particularly CTCF. The genome browser view visually confirms this indirect evidence: it not only shows co-localized ChIP-seq peaks for RAD21 and CTCF, but also highlights a distinct CTCF motif instance (the green block in **Figure 2F**) directly within the binding peak. By prioritizing this indirect evidence, the LLM Reasoning Agent inferred a medium-confidence prediction (score = 0.66) that matched the ChIP-seq peak. This example illustrates how Qimai leverages indirect evidence information to resolve ambiguous cases, enabling accurate prediction on non-TFs or novel proteins. Additional examples representing different scenarios are shown in **Supplementary Note E**.

## Discussion

The prediction of DNA-protein interactions remains a challenge in computational biology, particularly because existing models often fail to generalize to novel proteins or cases where DNA-binding is mediated indirectly. In this work, we introduce Qimai, an agent-based framework designed to address this challenge. Rather than relying on a single monolithic and ever-larger predictive model, Qimai intelligently synthesizes heterogeneous sources of information. Our results demonstrate that this new paradigm, using a Large Language Model (LLM) as a reasoning engine to integrate deep learning predictions with motif-based biological evidence, substantially improves generalization, particularly for proteins not seen during training.

The strategic choice of an LLM as the reasoning engine is central to Qimai’s success. Compared to a highly optimized LightGBM ensemble classifier trained on the same structured features, the LLM Agent achieved a 221% relative improvement in Matthews Correlation Coefficient (MCC), highlighting the value of flexible, context-aware reasoning over rigid, fix-weighted evidence of LightGBM (**Figure 2D**). As illustrated in the AP2A case study (**Supplementary Figure S2E**), Qimai correctly prioritized strong, reliable motif evidence over low-probability model prediction, which is similar to human expert’s reasoning. This performance gain is not restricted to a single proprietary model: open-source Gemma-3-27b-it performs comparably to proprietary Gemini models, particularly on the challenging “unseen” protein dataset (**Supplementary Figure S3B**). This confirms that the superior performance is rooted in the agent-based framework itself, which effectively leverages the general reasoning capacity of LLMs to synthesize disparate biological data into coherent, evidence-backed predictions.

A central biological insight from this study is the importance of modeling proteins that do not directly bind DNA. Many critical regulators, such as cohesin components (e.g., RAD21), chromatin remodelers, or enzymes like histone acetyltransferases, reach their target loci through recruitment by DNA-binding partners rather than by recognizing DNA sequences themselves. These proteins are indispensable for chromatin organization, enhancer–promoter communication, and transcriptional activation, yet they remain invisible to motif-centric models. Qimai’s indirect evidence pathway is specifically designed to capture these biologically important but challenging scenarios by considering motifs of known interaction partners and reasoning about plausible recruitment mechanisms. This capability to hypothesize how a protein is targeted to DNA, even in the absence of its own motif, represents a major conceptual advance over end-to-end black-box models.

The superior performance of Qimai particularly on unseen proteins arises from two deliberate design choices. First, foundation models like DNABERT and AlphaFold2 provide embeddings to our transformer DPI model with a more generalizable feature space than models trained on more specific sequence representations. Second, and more critically, the multi-agent reasoning layer incorporates direct and indirect motif evidence, creating a biologically grounded safety net for generalization. Ablation studies confirm that motif-based evidence is not a secondary input but a decisive factor that the LLM reasoning agent can elevate or discount depending on context. This design enables Qimai to generalize across diverse interaction models, direct, indirect, or structural, that underlie real regulatory systems.

Qimai’s modularity is another strength. The framework treats the predictive model as an interchangeable “Decision Agent.”, allowing seamless upgrades as new state-of-the-art architectures emerge. Our ablation studies on the seen proteins underscore this benefit (**Supplementary Figure S3C**). This adaptability ensures that the system can remain cutting-edge while preserving the interpretability and evidence-integration advantages of the agent-based structure.

There are limitations to our current implementation. Qimai depends on existing databases (CisBP, STRING) for motif and interaction data, which are themselves incomplete. As a result, the accuracy of predictions for non-DNA-binding proteins is partly constrained by the scope of existing annotations. Furthermore, our current approach emphasizes motif and sequence-derived features. A natural next step is to expand the Evidence Retriever to incorporate additional modalities such as epigenomic data (e.g., DNA methylation, histone modifications, chromatin accessibility), which provides crucial cell-type-specific context, as well as protein structure to assess the possibility of protein-protein interaction.

In conclusion, Qimai presents a new methodological paradigm for complex biological prediction tasks. By separating evidence gathering from evidence synthesis, and leveraging an LLM as the reasoning core, it creates a flexible and powerful framework that excels at generalization. This approach moves beyond simple black-box prediction, offering explainable, evidence-backed hypotheses that are more aligned with the process of scientific discovery. As foundation models and AI agents continue to evolve, frameworks like Qimai will be pivotal in translating vast, multi-modal biological data into actionable knowledge.

## Online Methods

### Data

We used a comprehensive set of human TF binding sites based on ChIP-seq experiments, where chromatin is immunoprecipitated by TF-specific antibody and the retained chromatin is then sequenced. The data (regions of enrichment also known as peak calls) we used in this study is generated by ENCODE with a uniform processing pipeline (**Supplementary Table 1**).

#### ChIP_690 dataset

This dataset contains 690 ChIP-seq experiments involving 158 TFs whose bindings were analyzed in different cell types or conditions. It was based on March 2012 ENCODE data freeze uniform TFBS peaks, which is a widely used dataset for training protein-DNA prediction models. Following *DeepBind* protocol^11^ for processing the data, positive samples are 512 base-pair (bp) DNA sequences centered on ChIP-seq binding peaks. The negative samples are shuffled positive sequences with matching dinucleotide composition. The shuffling was performed using the “fasta-dinucleotide-shuffle” package implemented in MEME^12^. For each experiment, 100 sequences (positive:negative=1:1) were randomly sampled. If the original set included less than 100 peaks, all sequences were kept. We randomly split the data into training/validation/test sets with a ratio of 7:1:2 (**Supplementary Table 2**). We used this dataset to benchmark the performance of the DPI model (this model is thus referred to as “DPI-bench”) with other models.

#### Encode3and4 dataset

This dataset contains 1877 ChIP-seq experiments involving 844 TFs, which includes all the available ChIP-seq experiments of phase III^13^ and phase IV of ENCODE by September 2023. Following *DeepSEA* protocol^4^ (see **Supplementary Note A** for this choice), human genome (hg38) was split into non-overlapping 200 bp bins. For each of the ChIP-seq experiments, we filtered out the binding peaks with signal value smaller than 50 and ran the intersection operation genome-wide with all 200 bp bins by constraining over a minimum overlapping of 50%. Then we kept the 200-bp bins with at least one TF binding event. Training samples are 512 bp DNA sequences centered at 200 bp bins. Some TFs have multiple experiments in different cell types or conditions. We took the majority vote as the final label if the training DNA sequence has different labels for the same TF. The bedtools software was used for genomic operations including intersection, merge, and overlap. Training/validation/test DNA sequences were split by chromosomes and strictly non overlapping. Each DNA-TF pair has a label of either 0 (not binding) or 1 (binding). We performed data down-sampling by sampling the negative DNA-TF pairs to keep the ratio of positive/negative samples as 1 (**Supplementary Table 3**). We used this dataset to train the final production model used in Qimai framework (referred to as “DPI”).

### DPI Model

DPI model is adapted from the original Transformer architecture in two aspects: (1) DNA as “source “language and protein as “target” language; (2) instead of next-word prediction task, DPI is to predict binary labels indicating interaction or not. Therefore, we removed the masking step in the decoder and exposed the full sequence to the model training. We described the model details (**Figure 1B; Supplementary Figure S1B**) in the following subsections.

#### Embedding extraction

We utilized a pre-trained DNA language model, DNABERT^6^, to transform DNA sequences into real-valued vectors. The pre-trained DNABERT-6mer model was selected since it achieves the best performance. The maximum length of DNA sequence for DNABERT is 512 bp. The hidden dimension of DNA embedding is 768. Protein sequences were downloaded from the UniProt database. AlphaFold2^14^ was applied to extract protein sequence embeddings. The maximum length of protein sequences for extraction is 1200 amino acids (AAs) due to computational limitations, which account for 95% of proteins in the dataset. The hidden dimension of protein embedding is 384.

#### Encode module

To capture the intra-protein and intra-DNA interactions, protein and DNA embeddings were processed in the Transformer encoder module separately. The encoder module consists of two sub-layers: the multi-head self-attention layer and fully connected feed-forward layer. A skip/residual connection structure and layer-wise normalization are incorporated around each sublayer to greatly facilitate training. The key feature within encoder is the multi-head self-attention layer which allows the model to associate all the relevant positions in a context to better encode a specific sequence and develop the “contextual understanding” in different aspects. DNA and protein encoders share the same architecture but not the learned weights. Inside of each encoder, the embedding is first projected into the hidden dimension of 240. Then the padded region is masked out in self-attention blocks. The output of encoder is refined embedding incorporating intra-sequence features.

#### Decoder module

To capture the interactions between DNA and protein, outputs of encoders will be sent to the decoder module, which adopts the original Transformer decoder architecture. Decoder module has similar architecture as encoder, except it calculates the cross-attention between DNA and protein instead of self-attention. Inside the decoder, interactions between two entities were learned and stored in the weights of cross attention matrices, which can be extracted later for interpretation. The output of decoder is interaction vector incorporating interaction information critical for binary prediction.

#### Prediction module

Lastly, we averaged the interaction vector across sequence dimensions to get a 240-dimensional vector. Then the vector goes through several linear layers with ReLU activations before the last linear layer, where a real-valued score is generated and sigmoid function is applied to get binary prediction.

### Implementation and training

DPI model is implemented in PyTorch 1.12.1 with CUDA 11.6 and trained on a NVIDIA A6000 with 48 GB of memory. It minimizes the binary cross-entropy loss with the Adam optimizer. While Adam does not require a learning rate scheduler, we observed adding a LambdaLR scheduler improved performance. Given the large number of hyperparameters, we did not perform a comprehensive grid search due to high computational cost. Instead, we identified several key hyperparameters based on early validation performance (**Supplementary Table 4**).

We found that the model showed better performance with increasing sequence length, which holds true for both protein and DNA. Restricted by DNABERT model constriction, we used the maximum length of 512-bp for DNA. Restricted by computational memory, we used the maximum length of 768 AAs for protein. If a sequence exceeds this length, we randomly select a continuous 768 AA-long subset for each training instance, ensuring varied exposure to different sequence regions. Because we trained using mini-batches, we zero-pad sequences shorter than 768 AAs.

The other hyperparameters were optimized using the Ray Tune framework for efficient exploration of the search space. Hidden dimension (hid_dim) is optimized in the range of (96, 240). Batch size is optimized in the range of (2, 20). Number of layers is optimized in the range of (1, 6). Number of heads is chosen between 6 and 12. Learning rate is optimized in the range of (5e-5, 0.5). The primary metric used for model performance evaluation were validation loss and validation accuracy. We trained two models for benchmark (DPI-bench) and production (DPI) runs respectively. It took 150 hours on a single A6000 GPU card with 48G memory to train the full model for production run (DPI) while 20 hours to train the DPI-bench model on the same hardware. The optimal hyperparameters for each training process were summarized in **Supplementary Table 4**.

### Interpretation of protein attention weights

We extracted the self-attention weights from protein encoder module. For each protein, the attention weights were averaged across all layers, heads and testing instances. We then identified regions with significantly large attention values. First, we calculated the mean and sd (standard deviation) of the whole sequence as background. Next, we calculated z-score for each value in the sequence and found sub-sequences no less than 8 AAs with z-scores larger than 0.2 (adjusted p-value of 0.001). Lastly, we combined the sub-sequences if they are within a specified range. We downloaded the DNA-binding domain information for all the TFs from the UniProt database and compared them with the regions identified before.

### CBenchmark with other methods

We compared the performance of DPI-bench with other popular TFBS prediction models such as DeepBind, DeepSEA, and DNABERT. The details of the training and dataset can be found in **Supplementary Notes** of original DNABERT^6^ publication.

#### Adaptation of other models for Qimai

To serve as alternative “Decision Agents” in the Qimai framework, we adapted both DeepSEA and Sei (the most updated version of DeepSEA with more complex convolutional neural network (CNN) architecture) models for our dual-input task. The original DNA-only CNN model was modified to accept a protein identifier as a second input. This identifier is mapped to a 50-dimensional embedding via an *nn.Embedding* layer, which is then concatenated with the flattened DNA feature vector before being passed to fully connected layers for final prediction. Training dataset is Encode3and4, which the same as that of DPI model (**Supplementary Table 3**). The training hyperparameters are listed in **Supplementary Table 4**. The performance on Encode3and4 test set for each of the prediction model is summarized in **Supplementary Table 5**. Sei performed the best in this test set which included unseen DNA sequences and seen proteins.

### Strategy for predicting unseen proteins with DeepSEA and Sei

To enhance the performance of DeepSEA and Sei, we employed the following strategy for predicting unseen protein:

1. **Prediction via sequence homology.** First, we check whether query protein has homologs (e.g. at least 40% sequence identity) in the model’s training set using the sequence aligner MMseqs2^15^. If homologs are found, an ensemble prediction is generated by running the model separately for each of the top N (e.g., N=3) most similar homologs and then averaging their output logits to produce a final, homology-informed probability.
2. **Prediction via shared TF family.** If no homolog is identified, we then check whether any training set protein belongs to the same TF family as the query using TF family defined in the UniProt database. If so, an ensemble prediction is generated by averaging the model’s output logits for these functionally related proteins.
3. **Generalized baseline prediction.** If neither sequence homology nor shared TF family provides a match, a generalized baseline prediction is made using the average embedding vector of all proteins in the training set.

#### Evidence Retriever Module

##### Direct Motif evidence

For the query protein, known binding motifs are fetched from a local instance of the CisBP database (downloaded on 2025.04.16). If multiple motifs exist, up to three are selected based on a priority score favoring directly determined motifs and those from reputable sources like JASPAR^16^ or HOCOMOCO^17^. The fetched Position Weight Matrices (PWMs) are converted to MEME format. The input DNA sequence is then scanned for these motifs using FIMO from the MEME suite, with a significance threshold of p-value (default: 1e-4). Results, including hit presence, p-values, scores, and CisBP motif reliability are summarized.

##### Indirect Motif evidence

The query protein is used to find potential interacting proteins from the STRING database^18^ for the Homo sapiens, filtered by a minimum interaction score of 700. Up to 50 interactors are considered. STRING interactors are cross-referenced with the CisBP TF list. For up to 3 known TFs (prioritized by STRING score), their motifs are fetched from CisBP as described above. The input DNA sequence is scanned for these indirect motifs using FIMO, and results are summarized similarly to direct motif evidence.

##### Prediction model-based evidence

We offered multiple deep learning models for probability prediction, primarily DPI, Sei, and DeepSEA. It is also possible to choose multiple models at the same time. Each model might have learned nuancedly different information, allowing for a more comprehensive interaction prediction.

#### Reasoning Agent LLM

The LLM receives the “Evidence Package” formatted into a specific prompt. Users can choose their preferred LLM, with options including Google’s Gemini-flash, Gemma3, or other powerful open-source models. The results shown here were generated using Gemini-2.0-flash. The core of our method lies in a dynamic prompting strategy which instructs the LLM how to weigh the evidence. For proteins that the computational model has seen before, the LLM would trust the model’s prediction more. For unseen proteins which are absent from model’s training set, the LLM would prioritize the biological motif evidence instead. Example prompts for unseen and seen proteins are shown in **Supplementary Figure S6**. After the LLM generates its analysis, the system automatically extracts the final prediction label and the reasoning.

#### Critique Agent & Final Output

The final component of the Qimai framework is a Critique Agent, implemented as a deterministic, rule-based system that assigns a final confidence score (ranging from 0 to 1) to the final prediction. This process provides a quantitative measure of certainty based on the coherence and strength of the underlying evidence. Starting from a baseline confidence value of 0.5, the confidence score is adjusted based on five criteria:

1. *Agreement/Disagreement*: Confidence is increased when the final prediction aligns with the presence of direct or indirect motifs and the predictions of the computational model(s). Conversely, confidence is penalized when the final prediction contradicts these evidence streams, with stronger penalties for contradicting high-quality evidence (e.g., direct motifs from reliable sources).
2. *Evidence Quality*: The magnitude of adjustments is weighted by the quality of the evidence. For example, bonuses are awarded for many motif hits or for hits with exceptionally significant p-values (e.g., <1e-7), while penalties are scaled back if motif scans had no hit or yielded ambiguous results.
3. *Protein Novelty*: The influence of each computational model’s prediction is modulated by a novelty factor. If a protein is unseen from the training set, the confidence (either positive or negative) from that computational model is reduced, reflecting a lower trust in its out-of-distribution performance.
4. *Prediction Model Certainty*: Independent of agreement, bonuses are awarded if a computational model’s prediction is highly confident (i.e., probability is very close to 0 or 1), as this suggests a clear signal from the model.
5. *LLM Explanation Quality*: The Qimai system performs a check for low-quality LLM outputs. If the explanation is unusually short, contains repetitive artifacts, or includes leftover template text, it is flagged as unreliable, and the confidence score is overridden to a near-zero value (e.g., 0.01).

A detailed summary of these heuristic rules and their default values is provided in **Supplementary Table 6.** The final output of Qimai is a triplet: the final prediction label (i.e. 0 for no binding and 1 for binding), the calculated confidence score, and the LLM’s textual explanation.

#### Implementation Details

The framework is implemented in Python 3, utilizing libraries such as LangGraph for workflow management, Pandas for data handling, PyTorch for deep learning models, Hugging Face Transformers for DNABERT model, BioPython for sequence manipulation, Requests for API calls (STRING DB), Google Generative AI SDK for Gemini, and DuckDB for local embedding databases. FIMO (MEME suite) is used as an external tool for motif scanning. Inference time varies across different LLM backends and prediction models with an average time of 20 seconds.

#### Benchmark Strategy and Performance Evaluation

We evaluated Qimai’s performance and generalization capabilities through a two-stage process: first, a rigorous internal cross-validation to assess robustness on held-out proteins from a single dataset of Encode3and4, and second, an external validation on a distinct dataset ChIP_690 enriched with novel proteins.

### Internal Robustness Assessment via Repeated Random Subsampling

To validate the robustness of our findings and ensure that Qimai’s generalization is not an artifact of a single train-test split, we performed a repeated random subsampling validation using Encode3and4 dataset. From the total pool of 844 proteins, we generated five independent splits. For each split, we randomly sampled 800 proteins to define the ‘seen’ training set for the underlying deep learning models. The remaining 44 proteins were held out as the ‘unseen’ test set for that specific partition.

To identify novel proteins in each held-out set, we performed a sequence-based analysis for each of the five splits. A reference sequence database was constructed from the 800-protein training set of that split. Each of the 44 held-out proteins was then queried against this database using MMseqs2. A protein was classified as “novel” for that split if its best hit failed to meet stringent homology criteria, defined as having a sequence identity of less than 30%, a bitscore below 50, or an E-value greater than 0.001. This process was repeated for all five splits, resulting in five distinct sets of novel proteins (ranging from 6 to 12 proteins per split) on which performance was specifically assessed (**Supplementary Figure S4A**).

### External Validation on a Novel Protein-Enriched Dataset

To further challenge the system’s generalization capabilities, we performed an external validation on the ChIP_690 dataset, which contains 78 “unseen” proteins that were not included in the Encode3and4 dataset. We analyzed whether these 78 proteins have homologs in the full 844-protein Encode3and4 training set using MMseqs2. A protein was classified as “novel” if its best hit failed to meet stringent homology criteria, defined as having a sequence identity of less than 30%, a bitscore below 50, or an E-value greater than 0.001. The ChIP_690 contains 51 novel proteins out of the 78 unseen proteins (65%).

### Evaluation and Ablation Setup

We evaluated the performance of the full Qimai system against the standalone predictive models (DPI, Sei, and DeepSEA) across two validation schemes. For our internal 5-fold cross-validation, we generated a balanced test set of 1000 examples involving 44 held-out proteins for each partition. For the external dataset assessment, we evaluated performance on a balanced set of 500 examples, which included all interactions involving the 78 previously unseen proteins. This external validation was repeated 10 times using 10 different random seeds, and the averaged performance with standard variation is reported.

Furthermore, to probe the contribution of different motif evidence streams, we conducted ablation studies where indirect motif information (“Qimai (w/o indirect)”) or direct motif information (“Qimai (w/o direct)”) was removed from the system. The system’s confidence scores were interpreted as probabilities of positive interaction; for negative interactions, the probability was taken as 1 minus the confidence score. These probabilities were used to generate Receiver Operating Characteristic (ROC) and Precision-Recall (PR) curves. A default probability threshold of 0.5 was used to derive binary classifications from the standalone predictive models, while the system’s binary prediction was parsed directly from the LLM’s response.

#### Evaluating the LLM Agent against a state-of-the-art Classifier

To rigorously evaluate the contribution of the LLM’s reasoning capabilities, we compared the performance of our LLM Agent against a state-of-the-art ensemble classifier. All evaluations were performed on a hold-out test set of 5,000 protein-DNA interaction pairs, each involving proteins that were unseen during the training of any underlying prediction models.

We chose LightGBM, a state-of-the-art gradient boosting model for comparison. The model was trained on a balanced dataset of 10,000 protein-DNA pairs (7,500 for training, 2,500 for internal validation and early stopping). To ensure a fair comparison, the baseline was provided with all the same structured evidence available to the LLM Agent. We collected a comprehensive set of 16 features designed to capture the nuances of the available evidence (**Supplementary Table 10**). These features were organized into three categories:

1. *Direct Motif Evidence*: Features describing the protein’s own binding motifs, including the number of significant hits, the −*log*10(*p*_*value*) of the best hit to represent statistical significance, and an ordinal score for the reliability of the motif source.
2. *Indirect Motif Evidence*: Features derived from the protein’s interaction partners, such as the number of interacting transcription factors, the maximum STRING-DB interaction score, and the −*log*10(*p* −*value*) of the best indirect hit.
3. *Deep Learning Model Predictions*: Features summarizing the outputs of the underlying deep learning models, including the raw binding probability, and a binary flag indicating if the protein was novel to that model’s training set.

Hyperparameters for the LightGBM model were selected to ensure robust generalization. We used a standard learning_rate of 0.05 and a num_leaves of 31 to balance model complexity and performance. The most critical hyperparameter, the number of boosting rounds (n_estimators), was determined automatically using early stopping with a patience of 10 rounds. This technique prevents overfitting by terminating training when the AUC score on the internal validation set ceased to improve, ensuring the model converged at its optimal iteration. The LightGBM model converged at an optimal iteration of 172.

The LightGBM model automatically performed feature selection, ultimately utilizing the 14 most informative features to make its predictions. Feature importances in LightGBM were calculated as the total gain, representing the cumulative reduction in loss contributed by each feature across all trees in the ensemble. Analysis of the trained model’s feature importances confirmed its ability to identify salient predictors (**Supplementary Table 10**). The final trained baseline achieved an AUC-ROC of 0.817 on its internal 2,500-sample test set, confirming it as a well-trained strong classifier. We assessed the performance of both the finalized LLM Agent and the trained LightGBM baseline on the 5,000 unseen test examples. Performance was measured using five metrics: Accuracy, F1-Score, Matthews Correlation Coefficient (MCC), AUC-ROC, and AUC-PR.

#### Evaluation on Qimai’s explanations

To systematically quantify the reasoning quality of the Qimai agent’s explanations, we implemented an ‘LLM-as-a-Judge’ framework. Three independent judge LLMs (open-source models phi4, deepseek-r1, qwen3 were used) evaluated each of the 500 explanations against its corresponding evidence package. Each judge generated scores on a 1-5 scale for the across five dimensions: Factual Accuracy, Evidence Coverage, Conflict Resolution, Clarity and Conciseness, and Overall Quality. Scores were collected in a structured format for quantitative analysis.

## Supplementary Tables

**Supplementary Table 1:** ChIP-seq experiment statistics.

| Dataset | Experiments | TFs |
| --- | --- | --- |
| ChIP_690 | 690 | 158 |
| Encode3and4 | 1877 | 844 |

**Supplementary Table 2:** Data statistics for DPI model used for benchmark.

| Split | Total pairs | Positive pairs | Negative pairs | # of proteins | # of DNAs |
| --- | --- | --- | --- | --- | --- |
| Training | 43,432 | 21,716 | 21,716 | 158 | 43,432 |
| Validation | 10,858 | 5,429 | 5,429 | 158 | 10,858 |
| Test | 21,546 | 10,773 | 10,773 | 158 | 21,546 |

**Supplementary Table 3:** Data statistics for DPI model used for production run.

| Split | Total pairs | Positive pairs | Negative pairs | # of proteins | # of DNAs |
| --- | --- | --- | --- | --- | --- |
| Training | 32,318,586 | 16,159,293 | 16,159,293 | 844 | 724,514 |
| Validation | 1,409,192 | 704,596 | 704,596 | 844 | 34,904 |
| Test | 2,567,990 | 1,283,995 | 1,283,995 | 844 | 65,180 |

**Supplementary Table 4:** Training hyperparameters for prediction models.

| Model | DPI-bench | DPI | DeepSEA | Sei |
| --- | --- | --- | --- | --- |
| max_dna_seq | 512 | 512 | 512 | 512 |
| max_protein_seq | 512 | 768 | - | - |
| protein_emb_dim | 384 | 384 | 50 | 50 |
| hid_dim | 240 | 240 | - | - |
| n_layers | 6 | 2 | - | - |
| n_heads | 12 | 6 | - | - |
| batch_size | 16 | 128 | 64 | 64 |
| epochs | 50 | 2 | 10 | 10 |
| learning_rate | 0.01 | 0.05 | 0.001 | 0.001 |

**Supplementary Table 5:** Prediction model performance on Encode3and4 test set.

| Model | Accuracy | F1<br>Score | MCC | Recall | Precision | Specificity | AUC-ROC | AUC-PR |
| --- | --- | --- | --- | --- | --- | --- | --- | --- |
| DPI | 0.820 | 0.822 | 0.640 | 0.837 | 0.812 | 0.803 | 0.900 | 0.887 |
| DeepSEA | 0.765 | 0.776 | 0.533 | 0.814 | 0.742 | 0.716 | 0.843 | 0.819 |
| Sei | 0.839 | 0.843 | 0.680 | 0.861 | 0.825 | 0.818 | 0.920 | 0.913 |

**Supplementary Table 6:** Heuristic rules and parameters for confidence score calculation.

| Category | Condition | Description | Seen<br>protein | Unseen<br>protein | Code variable name |
| --- | --- | --- | --- | --- | --- |
| Baseline | Initial value | Sets the initial score before adjustment | 0.50 |  | CONFIDENCE_BASELINE |
|  | Label=1 & Found | Bonus for agreement | +0.10 | +0.30 | DIRECT_MATCH_BOOST |
| **Direct Motif** | ... & Reliable | Additional bonus for high-quality evidence | +0.05 | +0.10 | DIRECT_RELIABLE_MATCH_EXTRA |
|  | Label=1 & Not Found | Penalty for missing evidence when predicting positive | -0.10 | -0.20 | DIRECT_MISSING_PENALTY |
|  | Label=0 & Found | Penalty for contradiction | -0.20 | -0.35 | DIRECT_MISMATCH_PENALTY |
|  | Label=0 & Not Found | Bonus for consistent absence of evidence | +0.05 | +0.15 | DIRECT_ABSENCE_BOOST |
| **Indirect Motif** | Label=1 & Found | Smaller bonus for indirect agreement | +0.10 | +0.20 | INDIRECT_MATCH_BOOST |
|  | Label=1 & Not Found | Smaller penalty for missing indirect evidence | -0.05 | -0.10 | INDIRECT_MISSING_PENALTY |
|  | Label=0 & Found | Smaller penalty for indirect contradiction | -0.15 | -0.25 | INDIRECT_MISMATCH_PENALTY |
|  | Label=0 & Not Found | Smaller bonus for consistent absence | +0.05 | +0.10 | INDIRECT_ABSENCE_BOOST |
| **Evidence Quality** | Num. Hits > 1 | Bonus per hit, up to a max value | +0.02/hit | +0.03/hit | ...NUM_HITS_SCALE_FACTOR |
|  | p-value is very low | Bonus for highly significant motif hits | +0.07 | +0.10 | ...LOW_PVALUE_BONUS |
|  | (Threshold) | p-value threshold for the above bonus | < 1e-7 | < 1e-8 | ...LOW_PVALUE_THRESHOLD |
| **Model Prediction** | Vote agrees w/ model | Bonus for model agreement | +0.30 | +0.10 | TRANSFORMER_MATCH_BOOST |
|  | ... & Confident | Add'l bonus for high-confidence agreement (prob > 0.8 or < 0.2) | +0.10 | +0.05 | TRANSFORMER_CONFIDENT_EXTRA |
|  | Vote disagrees | Penalty for model disagreement | -0.25 | -0.15 | TRANSFORMER_MISMATCH_PENALTY |
|  | ... & Confident | Add'l penalty for high-confidence disagreement | -0.15 | -0.05 | ...CONFIDENT_MISMATCH_EXTRA |
| **Multi-model** | 2nd+ model agrees | Diminishing bonus for subsequent agreeing models | 0.3x of base bonus |  | SUBSEQUENT_MODEL_AGREEMENT_FACTOR |
|  | 2nd+ model disagrees | Diminishing penalty for subsequent disagreeing models | 0.3x of base penalty |  | ...MISMATCH_PENALTY_FACTOR |
|  | Model/motif conflict | Amplifies penalty if >=2 models agree with LLM but contradict motifs | 1.2x of base penalty |  | MOTIF_CONFLICT_AMPLIFICATION_FACTOR |
| **Novelty** | Protein is novel | Multiplicative factor to reduce impact of a model's adjustment | 0.5x | 0.7x | ...CONFIDENT_MISMATCH_EXTRA |
| **Quality control** | LLM explanation fails | If explanation is too short, repetitive or has template text | Score overridden to 0.01 |  | Hard-coded logic |

**Supplementary Table 7:**
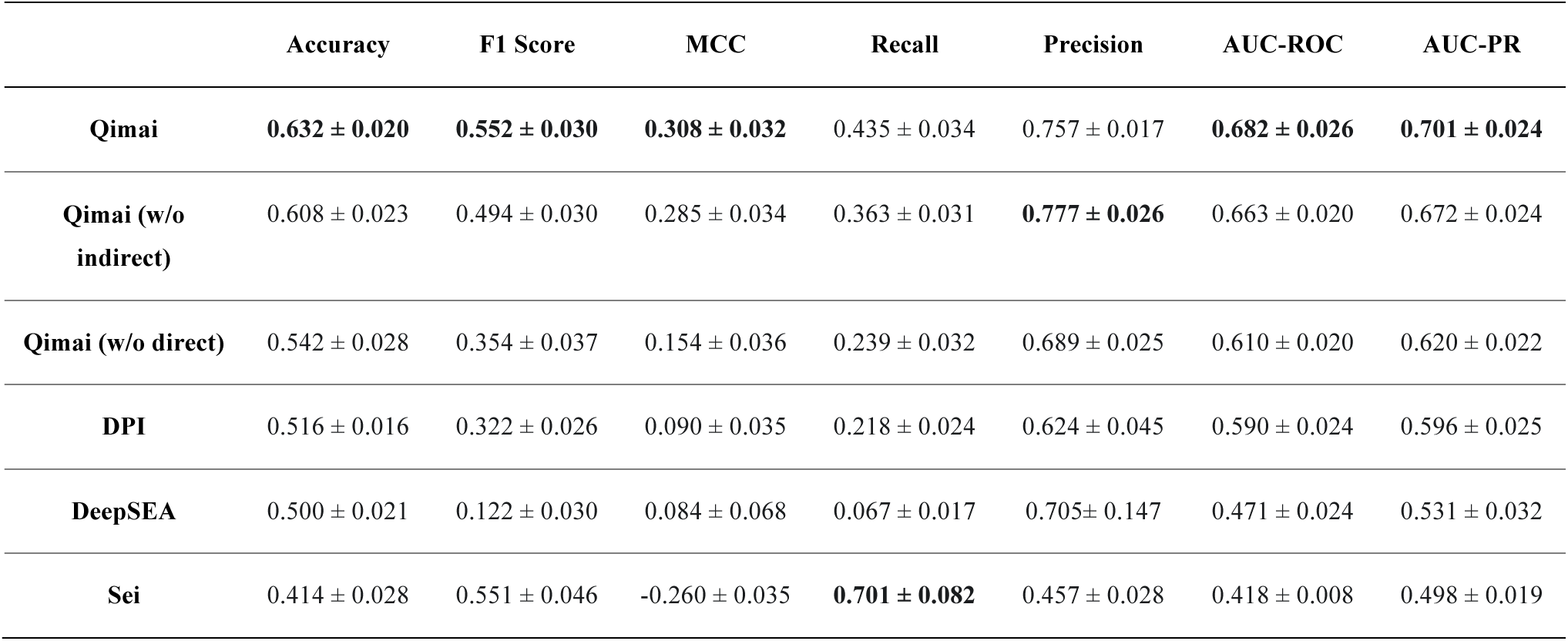
Performance summary on 78 unseen proteins with error bars representing the standard deviation over ten runs. Each run evaluated on 500 randomly sampled pairs.

|  | Accuracy | F1 Score | MCC | Recall | Precision | AUC-ROC | AUC-PR |
| --- | --- | --- | --- | --- | --- | --- | --- |
| **Qimai** | **0.632 ± 0.020** | **0.552 ± 0.030** | **0.308 ± 0.032** | 0.435 ± 0.034 | 0.757 ± 0.017 | **0.682 ± 0.026** | **0.701 ± 0.024** |
| **Qimai (w/o indirect)** | 0.608 ± 0.023 | 0.494 ± 0.030 | 0.285 ± 0.034 | 0.363 ± 0.031 | **0.777 ± 0.026** | 0.663 ± 0.020 | 0.672 ± 0.024 |
| **Qimai (w/o direct)** | 0.542 ± 0.028 | 0.354 ± 0.037 | 0.154 ± 0.036 | 0.239 ± 0.032 | 0.689 ± 0.025 | 0.610 ± 0.020 | 0.620 ± 0.022 |
| **DPI** | 0.516 ± 0.016 | 0.322 ± 0.026 | 0.090 ± 0.035 | 0.218 ± 0.024 | 0.624 ± 0.045 | 0.590 ± 0.024 | 0.596 ± 0.025 |
| **DeepSEA** | 0.500 ± 0.021 | 0.122 ± 0.030 | 0.084 ± 0.068 | 0.067 ± 0.017 | 0.705 ± 0.147 | 0.471 ± 0.024 | 0.531 ± 0.032 |
| **Sei** | 0.414 ± 0.028 | 0.551 ± 0.046 | -0.260 ± 0.035 | **0.701 ± 0.082** | 0.457 ± 0.028 | 0.418 ± 0.008 | 0.498 ± 0.019 |

**Supplementary Table 8:** Performance summary on 51 novel unseen proteins with error bars representing the standard deviation over ten runs.

|  | Accuracy | F1 Score | MCC | Recall | Precision | AUC-ROC | AUC-PR |
| --- | --- | --- | --- | --- | --- | --- | --- |
| **Qimai** | **0.653 ± 0.020** | 0.479 ± 0.038 | **0.314 ± 0.042** | 0.353 ± 0.037 | 0.759 ± 0.039 | **0.683 ± 0.035** | **0.650 ± 0.042** |
| **Qimai (w/o indirect)** | 0.621 ± 0.029 | 0.359 ± 0.061 | 0.273 ± 0.060 | 0.232 ± 0.048 | **0.812 ± 0.071** | 0.668 ± 0.034 | 0.636 ± 0.041 |
| **Qimai (w/o direct)** | 0.587 ± 0.030 | 0.337 ± 0.051 | 0.160 ± 0.053 | 0.231 ± 0.043 | 0.637 ± 0.056 | 0.628 ± 0.024 | 0.569 ± 0.036 |
| **DPI** | 0.570 ± 0.022 | 0.330 ± 0.044 | 0.122 ± 0.045 | 0.230 ± 0.038 | 0.593 ± 0.049 | 0.624 ± 0.022 | 0.569 ± 0.035 |
| **DeepSEA** | 0.585 ± 0.025 | 0.222 ± 0.049 | 0.166 ± 0.081 | 0.131 ± 0.030 | 0.731 ± 0.136 | 0.578 ± 0.051 | 0.536 ± 0.060 |
| **Sei** | 0.417 ± 0.034 | **0.580 ± 0.030** | -0.176 ± 0.099 | **0.890 ± 0.044** | 0.431 ± 0.025 | 0.489 ± 0.018 | 0.485 ± 0.028 |

**Supplementary Table 9:**
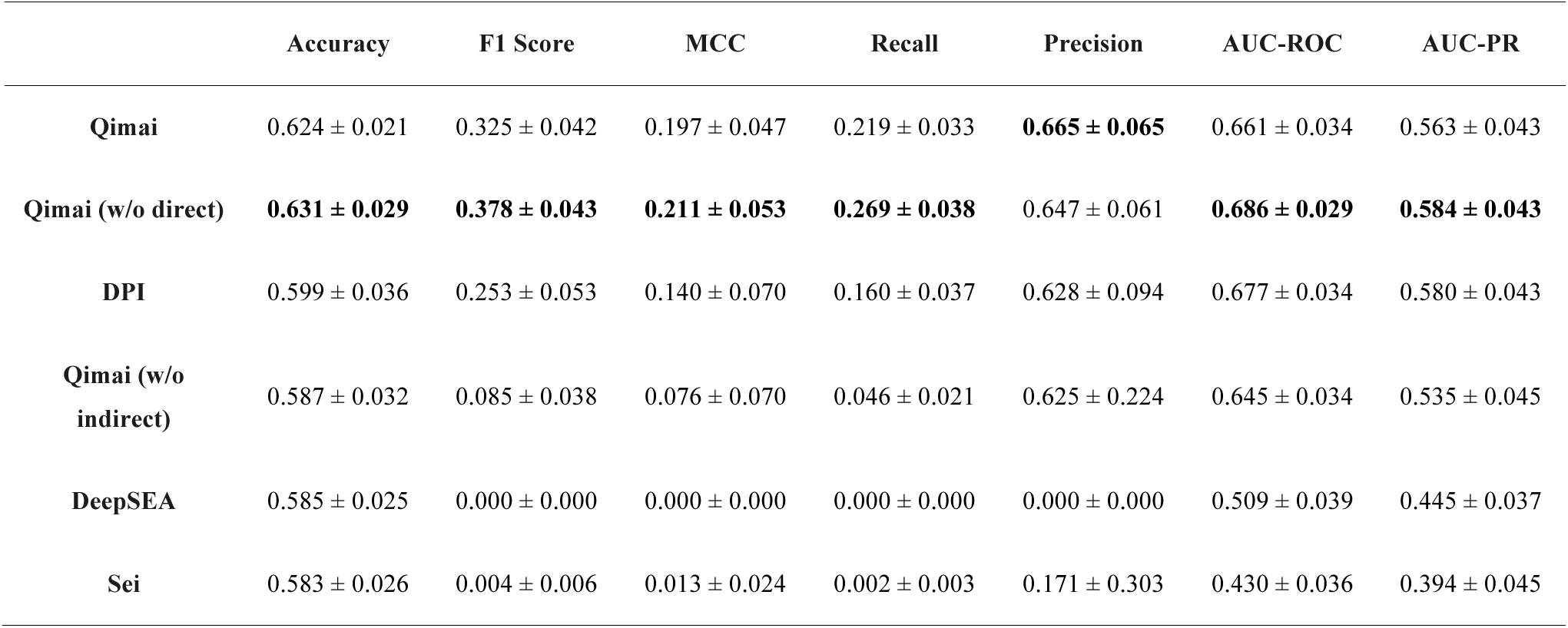
Performance summary on 45 novel proteins without direct motifs with error bars representing the standard deviation over ten runs.

|  | Accuracy | F1 Score | MCC | Recall | Precision | AUC-ROC | AUC-PR |
| --- | --- | --- | --- | --- | --- | --- | --- |
| Qimai | 0.624 ± 0.021 | 0.325 ± 0.042 | 0.197 ± 0.047 | 0.219 ± 0.033 | **0.665 ± 0.065** | 0.661 ± 0.034 | 0.563 ± 0.043 |
| Qimai (w/o direct) | **0.631 ± 0.029** | **0.378 ± 0.043** | **0.211 ± 0.053** | **0.269 ± 0.038** | 0.647 ± 0.061 | **0.686 ± 0.029** | **0.584 ± 0.043** |
| DPI | 0.599 ± 0.036 | 0.253 ± 0.053 | 0.140 ± 0.070 | 0.160 ± 0.037 | 0.628 ± 0.094 | 0.677 ± 0.034 | 0.580 ± 0.043 |
| Qimai (w/o indirect) | 0.587 ± 0.032 | 0.085 ± 0.038 | 0.076 ± 0.070 | 0.046 ± 0.021 | 0.625 ± 0.224 | 0.645 ± 0.034 | 0.535 ± 0.045 |
| DeepSEA | 0.585 ± 0.025 | 0.000 ± 0.000 | 0.000 ± 0.000 | 0.000 ± 0.000 | 0.000 ± 0.000 | 0.509 ± 0.039 | 0.445 ± 0.037 |
| Sei | 0.583 ± 0.026 | 0.004 ± 0.006 | 0.013 ± 0.024 | 0.002 ± 0.003 | 0.171 ± 0.303 | 0.430 ± 0.036 | 0.394 ± 0.045 |

**Supplementary Table 10:**
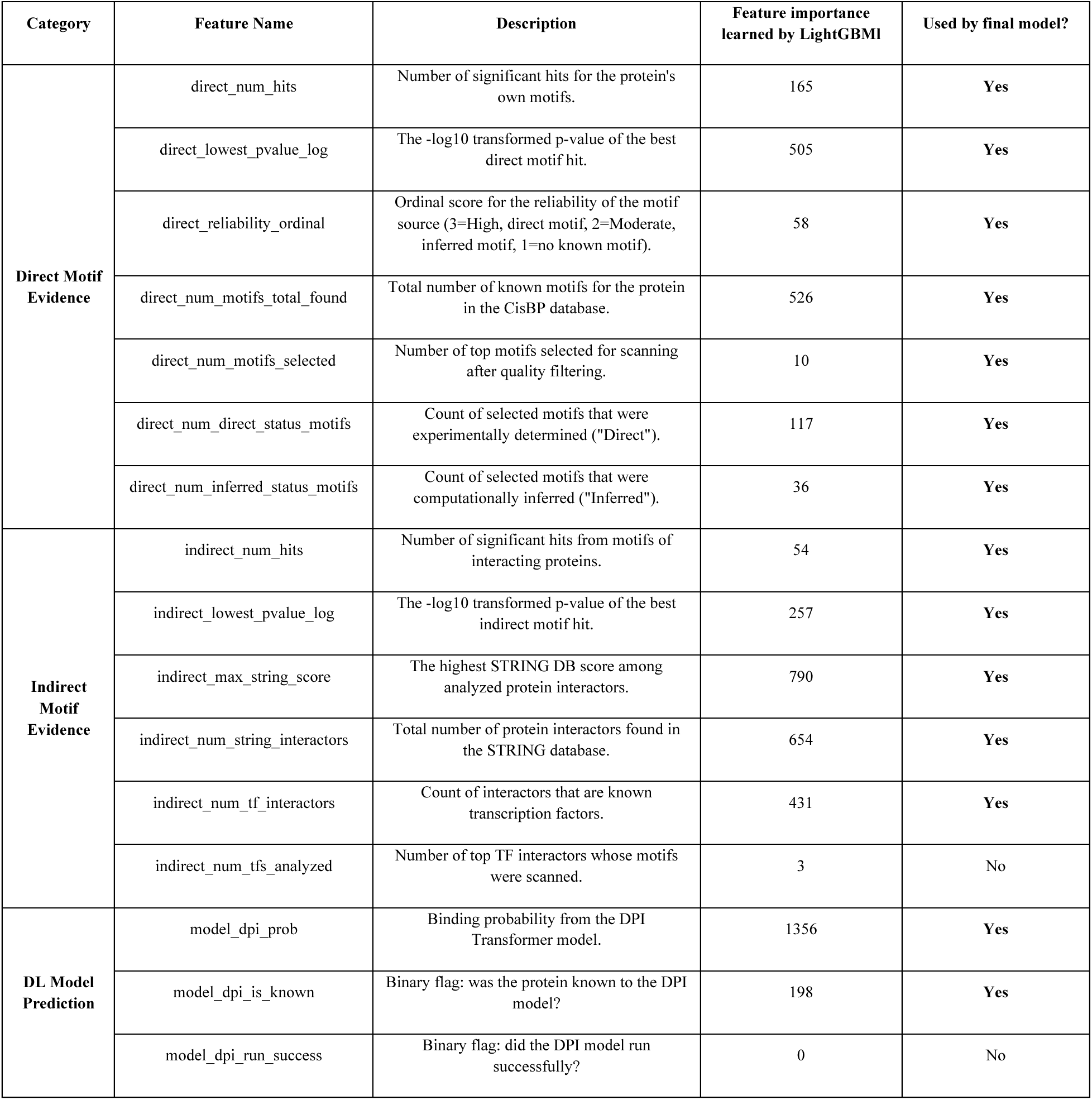
Features provided to the baseline LightGBM classifier.

**Supplementary Table 11:** Other variations of heuristic parameters for confidence score calculation.

| Category | Condition | Description | Default | V1 | V2 | V3 | Code variable name |
| --- | --- | --- | --- | --- | --- | --- | --- |
| **Baseline** | Initial value | Sets the initial score before adjustment | 0.50 | 0.50 | 0.45 | 0.55 | CONFIDENCE_BASELINE |
| **Direct Motif** | Label=1 & Found | Bonus for agreement | +0.30 | +0.25 | +0.20 | +0.28 | DIRECT_MATCH_BOOST |
|  | ... & Reliable | Additional bonus for high-quality evidence | +0.10 | +0.10 | +0.08 | +0.12 | DIRECT_RELIABLE_MATCH_EXTRA |
|  | Label=1 & Not Found | Penalty for missing evidence when predicting positive | -0.20 | -0.15 | -0.18 | -0.10 | DIRECT_MISSING_PENALTY |
|  | Label=0 & Found | Penalty for contradiction | -0.35 | -0.30 | -0.35 | -0.25 | DIRECT_MISMATCH_PENALTY |
|  | Label=0 & Not Found | Bonus for consistent absence of evidence | +0.15 | +0.10 | +0.08 | +0.15 | DIRECT_ABSENCE_BOOST |
| **Indirect Motif** | Label=1 & Found | Smaller bonus for indirect agreement | +0.20 | +0.10 | +0.08 | +0.15 | INDIRECT_MATCH_BOOST |
|  | Label=1 & Not Found | Smaller penalty for missing indirect evidence | -0.10 | -0.05 | -0.07 | -0.03 | INDIRECT_MISSING_PENALTY |
|  | Label=0 & Found | Smaller penalty for indirect contradiction | -0.25 | -0.15 | -0.12 | -0.10 | INDIRECT_MISMATCH_PENALTY |
|  | Label=0 & Not Found | Smaller bonus for consistent absence | +0.10 | +0.05 | +0.04 | +0.08 | INDIRECT_ABSENCE_BOOST |
| **Evidence Quality** | Num. Hits > 1 | Bonus per hit, up to a max value | +0.03/hit | +0.02/hit | +0.015/hit | +0.025/hit | ...NUM_HITS_SCALE_FACTOR |
|  | p-value is very low | Bonus for highly significant motif hits | +0.10 | +0.07 | +0.05 | +0.08 | ...LOW_PVALUE_BONUS |
|  | (Threshold) | p-value threshold for the above bonus | < 1e-8 | < 1e-7 | < 1e-8 | < 5e-7 | ...LOW_PVALUE_THRESHOLD |
| **Model Prediction** | Vote agrees w/ model | Bonus for model agreement | +0.10 | +0.15 | +0.12 | +0.20 | TRANSFORMER_MATCH_BOOST |
|  | ... & Confident | Add'l bonus for high-confidence agreement (prob > 0.8 or < 0.2) | +0.05 | +0.10 | +0.08 | +0.12 | TRANSFORMER_CONFIDENT_EXTRA |
|  | Vote disagrees | Penalty for model disagreement | -0.15 | -0.20 | -0.18 | -0.15 | TRANSFORMER_MISMATCH_PENALTY |
|  | ... & Confident | Add'l penalty for high-confidence disagreement | -0.05 | -0.10 | -0.12 | -0.08 | ...CONFIDENT_MISMATCH_EXTRA |
| **Multi-model** | 2nd+ model agrees | Diminishing bonus for subsequent agreeing models | 0.3x of base bonus |  |  |  | SUBSEQUENT_MODEL_AGREEMENT_FACTOR |
|  | 2nd+ model disagrees | Diminishing penalty for subsequent disagreeing models | 0.3x of base penalty |  |  |  | ...MISMATCH_PENALTY_FACTOR |
|  | Model/motif conflict | Amplifies penalty if >=2 models agree with LLM but contradict motifs | 1.2x of base penalty |  |  |  | MOTIF_CONFLICT_AMPLIFICATION_FACTOR |
| Novelty | Protein is novel | Multiplicative factor to reduce impact of a model's adjustment | 0.7x | 0.6x | 0.5x | 0.8x | ...CONFIDENT_MISMATCH_EXTRA |
| Quality control | LLM explanation fails | If explanation is too short, repetitive or has template text | Score overridden to 0.01 |  |  |  | Hard-coded logic |

**Supplementary Table 12:** Performance summary on 44 unseen proteins with error bars representing the standard deviation over five splits. Each run evaluated on 1000 randomly sampled pairs.

|  | Accuracy | F1 Score | MCC | Recall | Precision | AUC-ROC | AUC-PR |
| --- | --- | --- | --- | --- | --- | --- | --- |
| Qimai | **0.565 ± 0.017** | 0.537 ± 0.012 | **0.178 ± 0.018** | 0.440 ± 0.010 | 0.691 ± 0.031 | 0.632 ± 0.028 | **0.701 ± 0.036** |
| Qimai (w/o indirect) | 0.548 ± 0.026 | 0.501 ± 0.024 | 0.155 ± 0.032 | 0.396 ± 0.028 | 0.683 ± 0.027 | 0.618 ± 0.032 | 0.685 ± 0.036 |
| Qimai (w/o direct) | 0.516 ± 0.023 | 0.368 ± 0.038 | 0.156 ± 0.039 | 0.247 ± 0.028 | **0.730 ± 0.065** | **0.643 ± 0.029** | 0.693 ± 0.037 |
| DPI | 0.550 ± 0.042 | 0.399 ± 0.091 | 0.158 ± 0.088 | 0.287 ± 0.079 | 0.673 ± 0.091 | 0.612 ± 0.048 | 0.653 ± 0.065 |
| DeepSEA | 0.462 ± 0.042 | 0.152 ± 0.118 | 0.079 ± 0.107 | 0.089 ± 0.071 | 0.577 ± 0.368 | 0.528 ± 0.037 | 0.611 ± 0.057 |
| Sei | 0.562 ± 0.042 | **0.696 ± 0.040** | 0.018 ± 0.070 | **0.878 ± 0.037** | 0.577 ± 0.045 | 0.525 ± 0.021 | 0.601 ± 0.044 |

**Supplementary Table 13:** Performance summary on 6-12 novel held-out proteins with error bars representing the standard deviation over five splits.

|  | Accuracy | F1 Score | MCC | Recall | Precision | AUC-ROC | AUC-PR |
| --- | --- | --- | --- | --- | --- | --- | --- |
| Qimai | 0.693 ± 0.046 | 0.497 ± 0.234 | 0.272 ± 0.223 | 0.444 ± 0.230 | 0.577 ± 0.255 | **0.633 ± 0.160** | **0.593 ± 0.231** |
| Qimai (w/o indirect) | **0.700 ± 0.062** | 0.515 ± 0.242 | **0.290 ± 0.245** | 0.464 ± 0.230 | **0.585 ± 0.266** | 0.625 ± 0.159 | 0.573 ± 0.219 |
| Qimai (w/o direct) | 0.643 ± 0.065 | 0.348 ± 0.240 | 0.137 ± 0.236 | 0.277 ± 0.203 | 0.501 ± 0.295 | 0.588 ± 0.128 | 0.497 ± 0.200 |
| DPI | 0.620 ± 0.090 | 0.343 ± 0.200 | 0.114 ± 0.237 | 0.271 ± 0.162 | 0.498 ± 0.296 | 0.575 ± 0.113 | 0.502 ± 0.217 |
| DeepSEA | 0.608 ± 0.118 | 0.041 ± 0.092 | -0.020 ± 0.045 | 0.027 ± 0.061 | 0.084 ± 0.189 | 0.555 ± 0.082 | 0.445 ± 0.077 |
| Sei | 0.417 ± 0.115 | **0.560 ± 0.117** | 0.104 ± 0.068 | **0.985 ± 0.018** | 0.399 ± 0.116 | 0.514 ± 0.034 | 0.398 ± 0.106 |

## Acknowledgement

This work is partially supported by NIH (R01HG009626 to WW).

## Code availability

Source code used to generate all results shown in this paper is deposited and publicly available at https://github.com/Wang-lab-UCSD/DPI-agent.

## Supplementary Figures

**Supplemental Figure 1:**
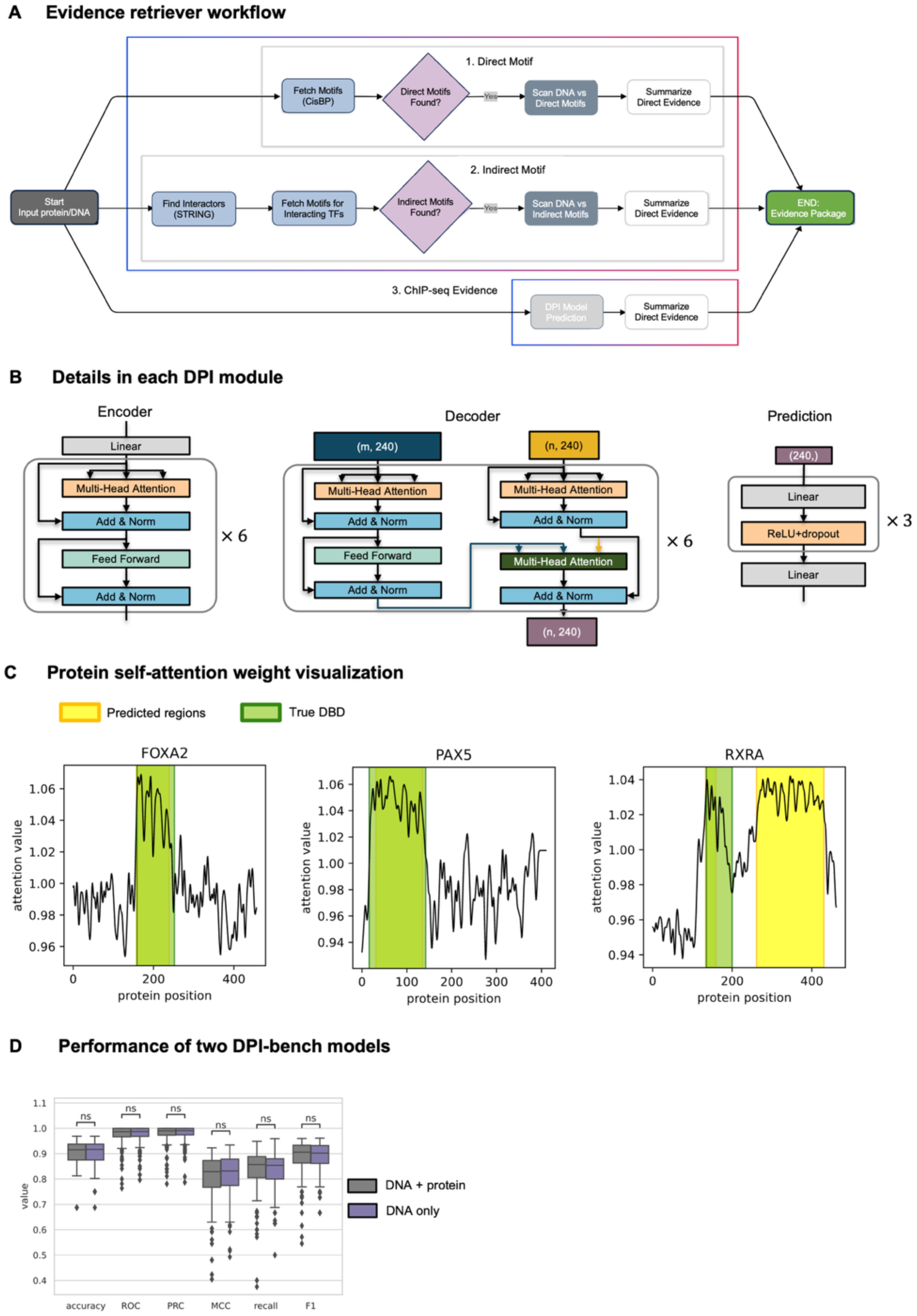
Framework details. **(A) Evidence Retriever workflow.** This module gathers three types of evidence: 1. Direct Motif evidence (fetching the query protein’s motifs from CisBP database and scanning the DNA); 2. Indirect Motif evidence (identifying protein interactors via STRING database, fetching their motifs, and scanning the DNA); 3. Computational Model Evidence (predicting interaction using a pre-trained model like the DPI transformer detailed in panel B, DeepSEA, or Sei). **(B) Details in each DPI module.** Encoder includes a linear transformation and the basic building block including two sub-layers: the multi-head self-attention layer and fully connected feed-forward layer. Decoder module has similar architecture as encoder, except it calculates the cross-attention between DNA and protein instead of self-attention. Prediction module includes linear layer with ReLU activation. **(C) Self-attention weights of protein sequences.** The regions with high attention weights extracted from encoder module are highlighted in yellow. True DNA-binding domains (DBDs) are highlighted in green. Yellow and green regions are overlapped. **(D) Performance of two DPI-bench models**, one utilizing both protein and DNA information (gray) and the other using DNA information only (purple). The two models showed similar performance. “ns” stands for not significant difference for t-test.

**Supplemental Figure 2:**
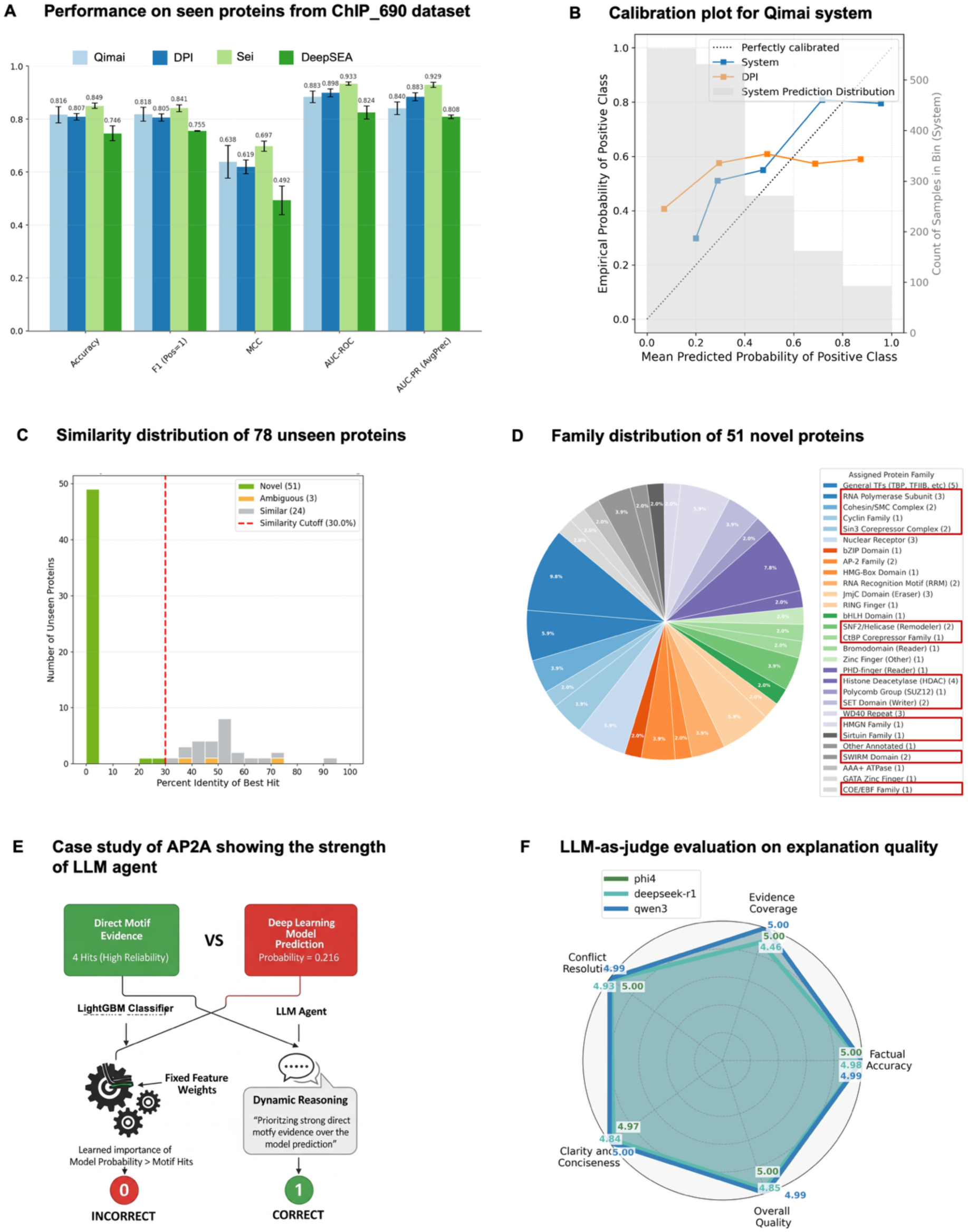
Qimai framework superior performance in external ChIP_690 dataset. **(A) Performance on the seen protein in ChIP_690 dataset**. The full Qimai system, incorporating the DPI model prediction and all motif evidence, is compared against standalone deep learning models (DPI, Sei, DeepSEA) on unseen DNA sequences with known proteins. The standalone Sei model shows the strongest performance, while the Qimai system is highly competitive. **(B) Calibration plot for the final System and standalone DPI model scores**. This plot compares the mean predicted probability of a positive interaction (x-axis) against the empirical fraction of actual positive interaction (y-axis). The System (blue) is well-calibrated, closely tracking the ideal diagonal reference line. In contrast, the DPI model (orange) is consistently over-confident. The background histogram displays the distribution of the System’s prediction scores, highlighting that its calibration is most effective in the highly populated low-confidence region. **(C) Similarity distribution of 78 unseen proteins**. “Novel Proteins” represent proteins from the unseen set that showed no significant sequence similarity to any protein in the training set, i.e. <30% sequence identity, a bitscore below 50, or an E-value greater than 0.001. “Ambiguous Proteins” are those unseen proteins that didn’t meet all the stringent criteria. “Similar Proteins” are those unseen proteins that did have at least one significant sequence match within the training set. The annotation is used throughout the paper. **(D) Family distribution of 51 novel proteins**. Families with red boxes are the novel families which are absent from training data. Almost half of the novel proteins also belong to the novel protein families (23/51). **(E) The LLM Agent’s context-aware reasoning resolves conflicting evidence where a state-of-the-art ensemble classifier fails**. This case study of protein AP2A illustrates a scenario with contradictory inputs: strong, high-reliability direct motif evidence (4 hits) versus a low-probability deep learning model prediction (0.216). The baseline classifier, constrained by its fixed, learned feature weights where model probability has a higher learned importance, is misled by the low score and makes an incorrect prediction. In contrast, the LLM Agent dynamically reasons about the evidence quality, explicitly prioritizing the strong motif data over the single computational score to arrive at the correct classification. **(F) LLM-as-judge evaluation of explanation quality**. A radar chart displaying the assessment of Qimai’s generated explanations by an LLM-as-judge. The explanations received near-perfect scores (out of 5) across five criteria, highlighting their factual accuracy, clarity, and effectiveness in resolving conflicting evidence.

**Supplemental Figure 3:**
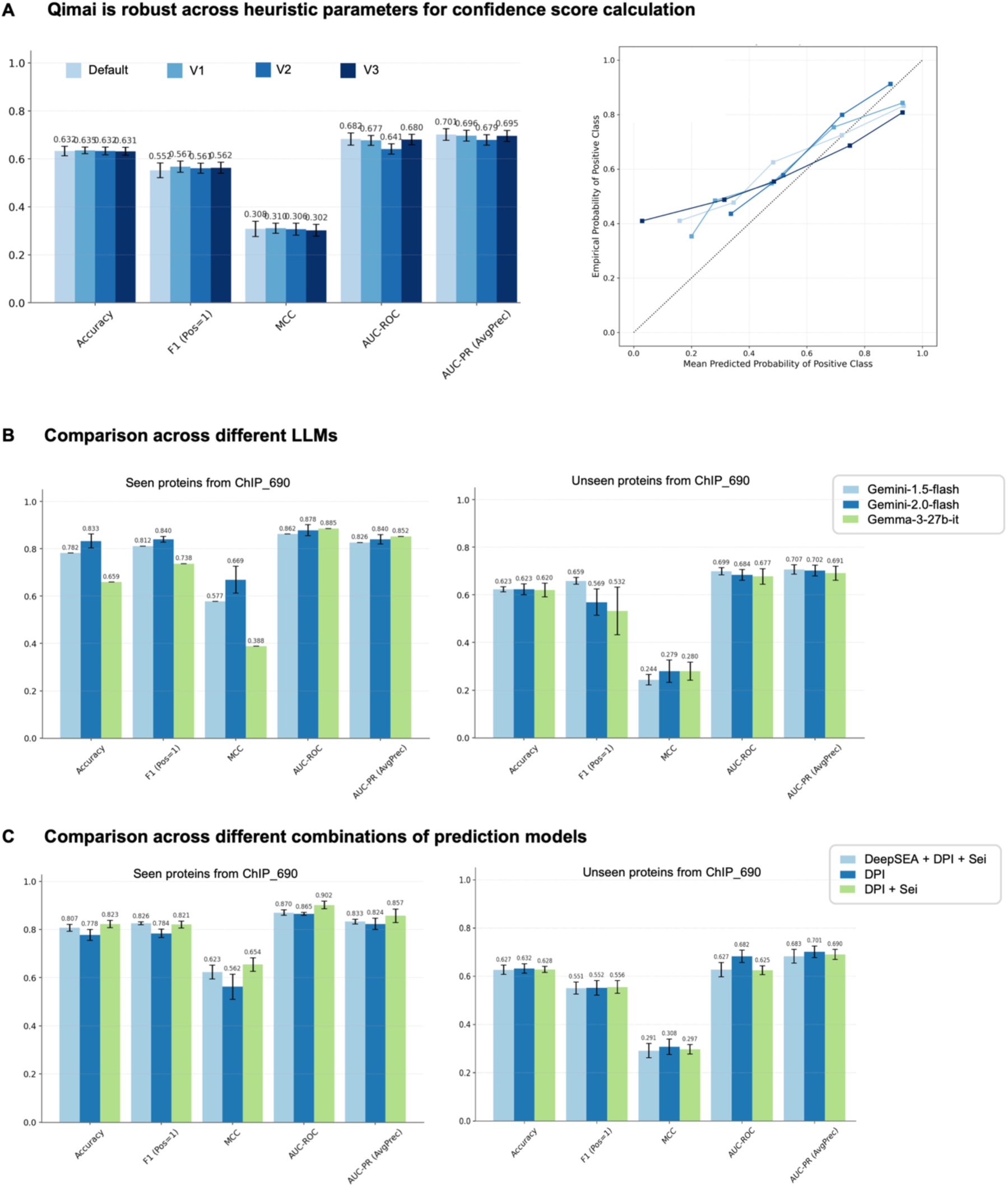
Evaluation of Qimai framework robustness in external ChIP_690 dataset. **(A) Qimai’s confidence score is robust to variations in heuristic parameter sets**. Left panel shows key metrics for the system’s predictions across four different heuristic configurations (**Supplementary Table 11**). The results demonstrate high and stable performance, with minimal variation across all test configurations. Error bars representing the standard deviation over ten runs. Each run was evaluated on 500 randomly sampled pairs. Right panel is the corresponding calibration curves. The close alignment of all four colored lines to this diagonal indicates that the confidence score remains a reliable and well-calibrated measure of prediction accuracy, regardless of the underlying heuristic model. **(B) Comparison across different LLM reasoning agents**. Qimai’s performance was evaluated using gemini-1.5-flash, gemini-2.0-flash, and the open-source gemma-3-27b-it on both the seen (left) and “unseen” (right) proteins in ChIP_690 dataset. Performance is shown to be robust and highly comparable across all LLMs, particularly on the challenging “unseen” dataset, demonstrating the framework’s consistency. Error bars in all panels represent the standard deviation over ten runs. Each run was evaluated on 500 randomly sampled pairs. **(C) Comparison across different combinations of prediction models**. Qimai’s performance was evaluated using three different combinations of prediction models. On the seen proteins in external dataset (left), integrating predictions from two models (“DPI + Sei”) yields the highest performance. In contrast, on the unseen proteins (right), providing only the single best-generalizing model (DPI) is the most effective strategy. Error bars in all panels represent the standard deviation over ten runs. Each run was evaluated on 500 randomly sampled pairs.

**Supplemental Figure 4:**
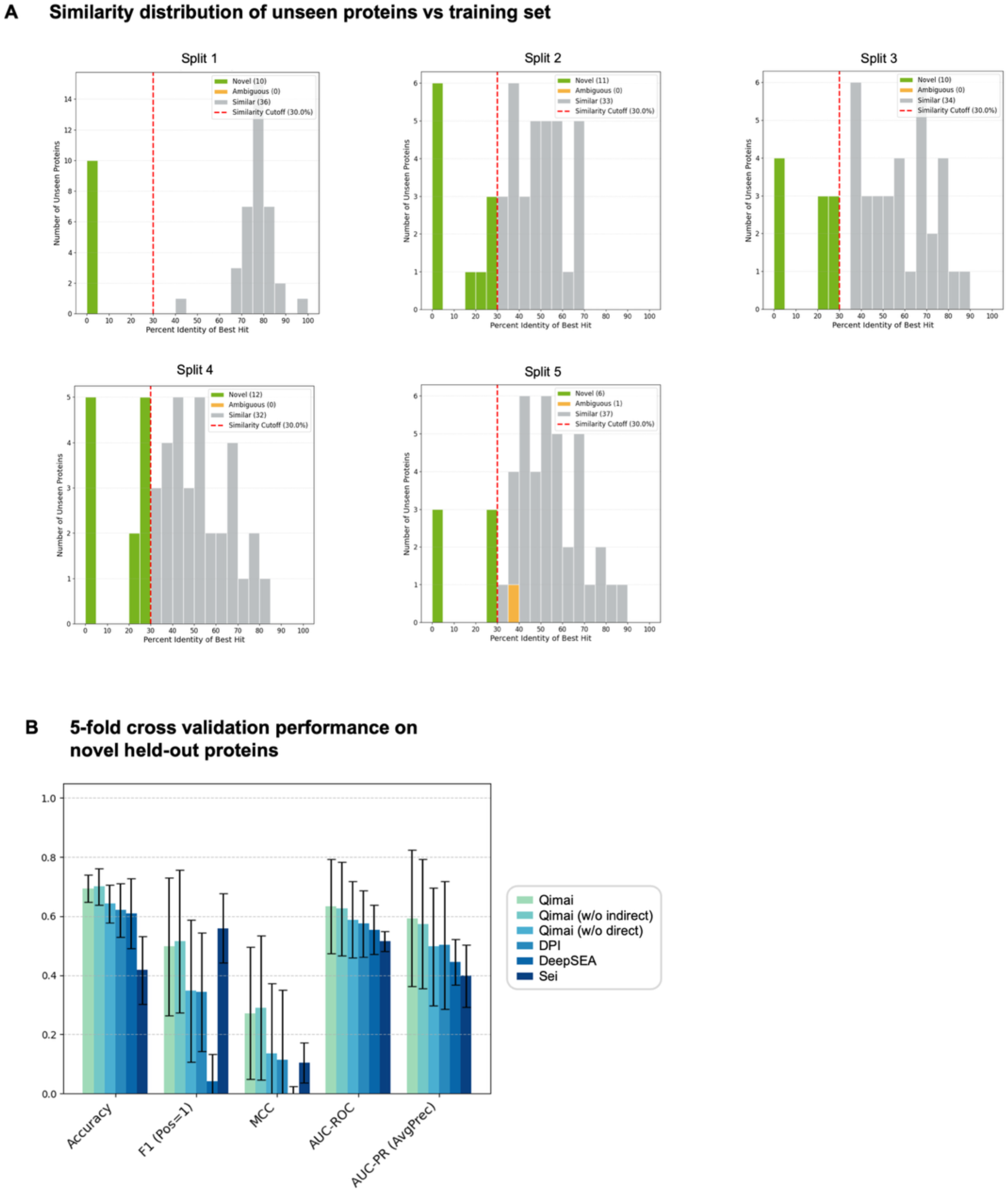
Internal Cross-Validation of Qimai performance. **(A) Similarity distribution of unseen proteins vs training set for each of the five splits**. “Novel Proteins” represent proteins from the unseen set that showed no significant sequence similarity to any protein in the training set, i.e. <30% sequence identity. “Similar Proteins” are those unseen proteins that had at least one sequence match within the training set. **(B) Performance across five random splits on novel proteins**. A comparison of the full Qimai System against standalone deep learning modles (DPI, Sei, DeepSEA) and its ablated version (“Qimai (w/o indirect)”, “Qimai (w/o direct)”) across five random splits. Results are averaged over five random splits with error bars showing standard deviation.

**Supplemental Figure 5:**
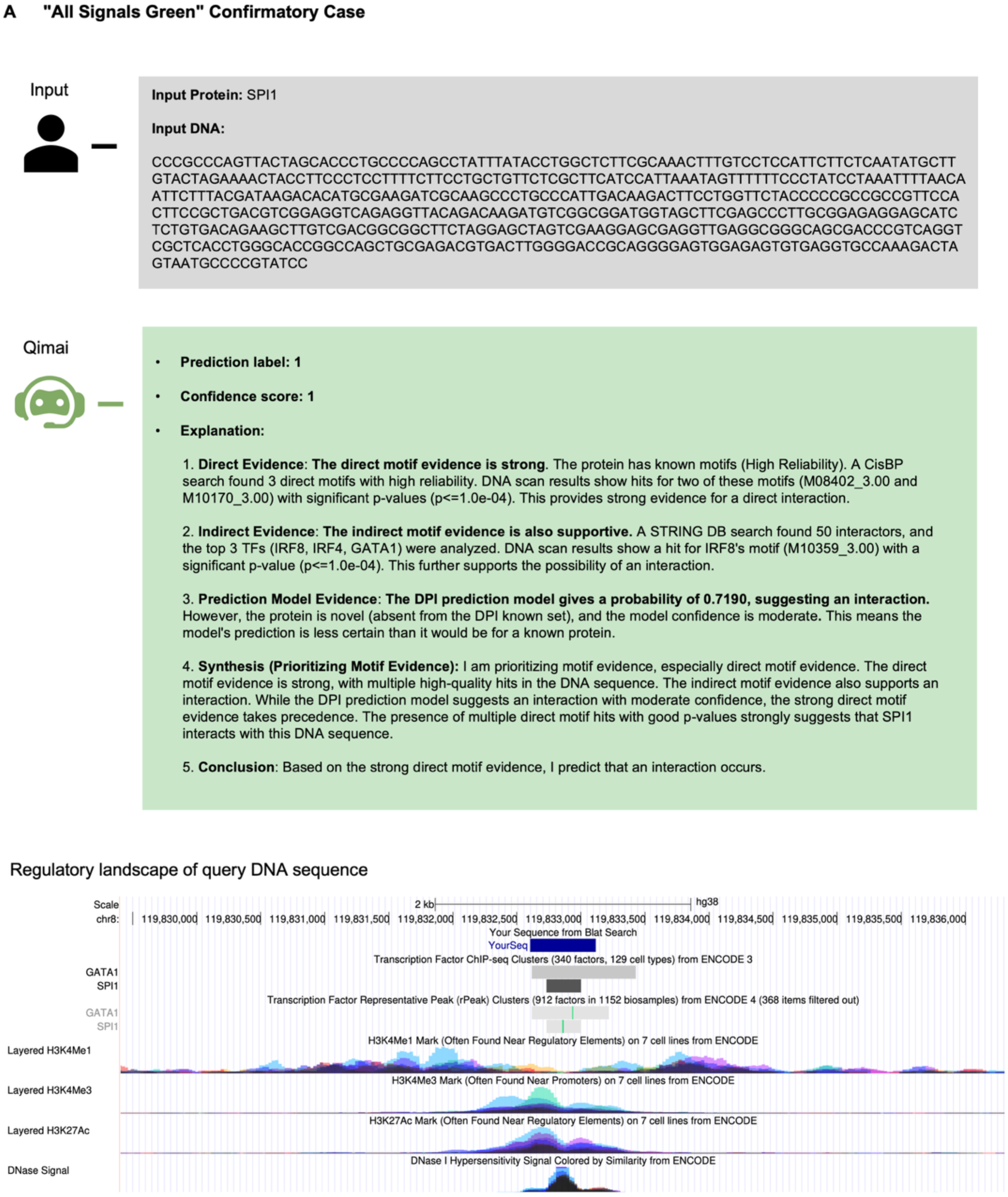
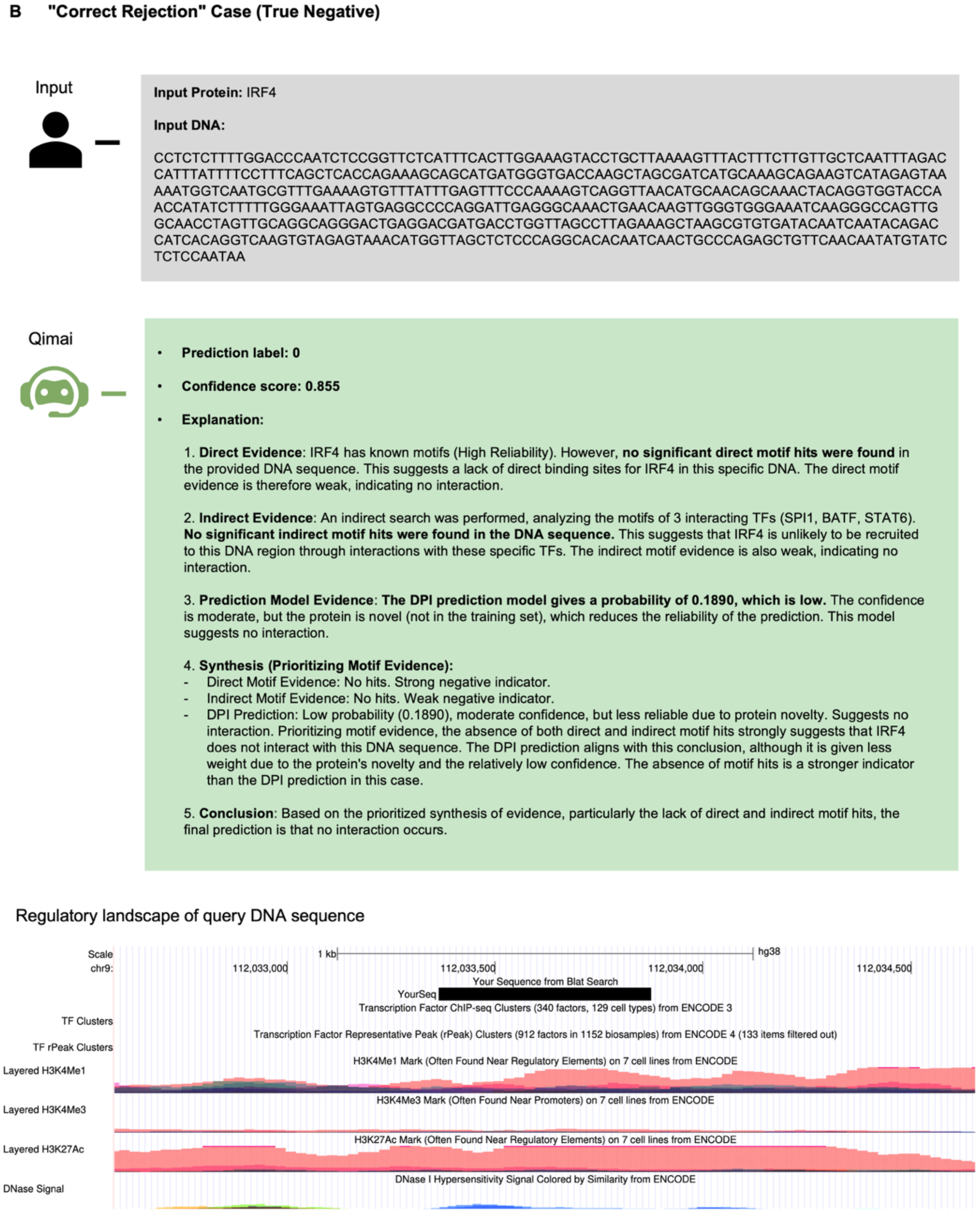
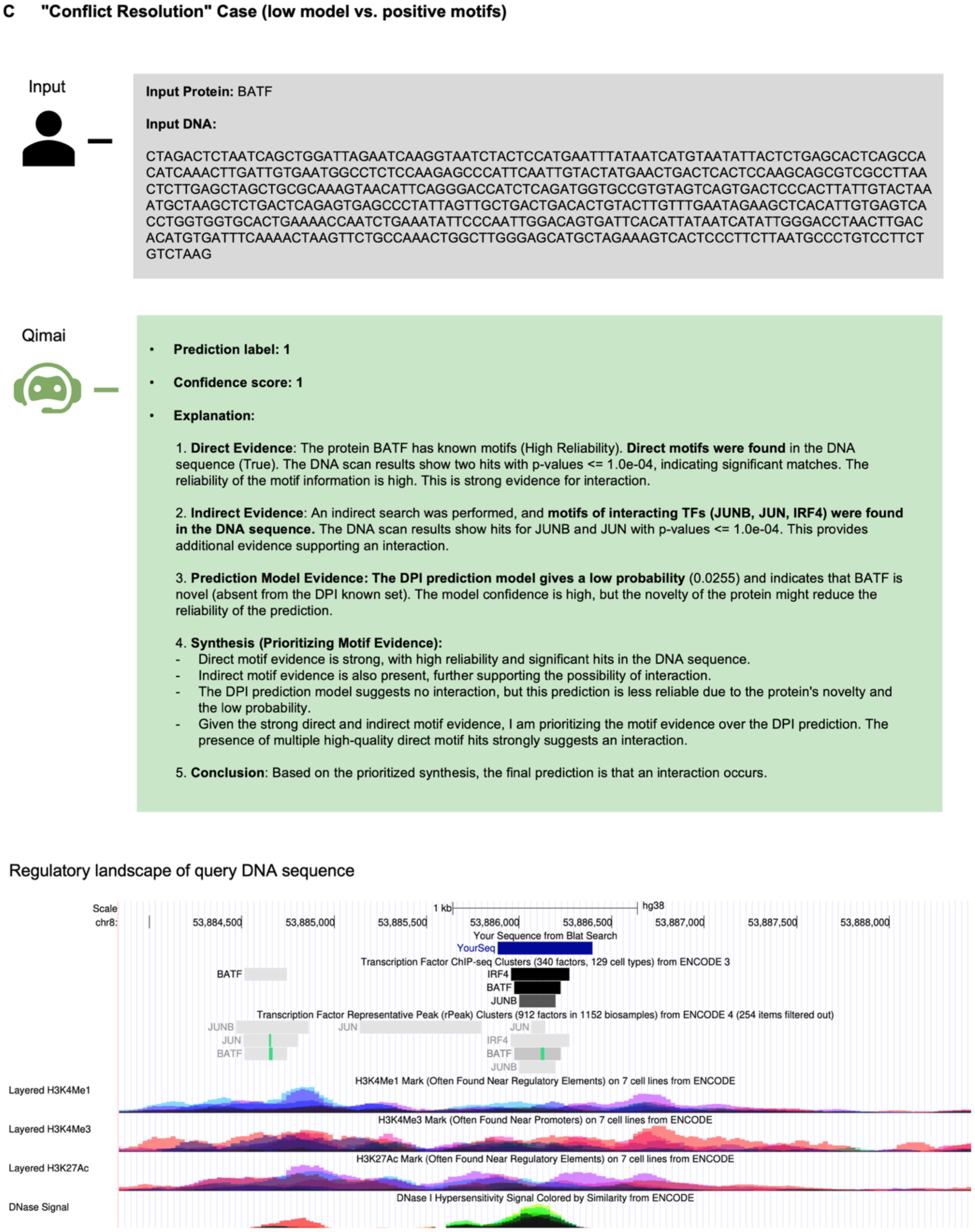
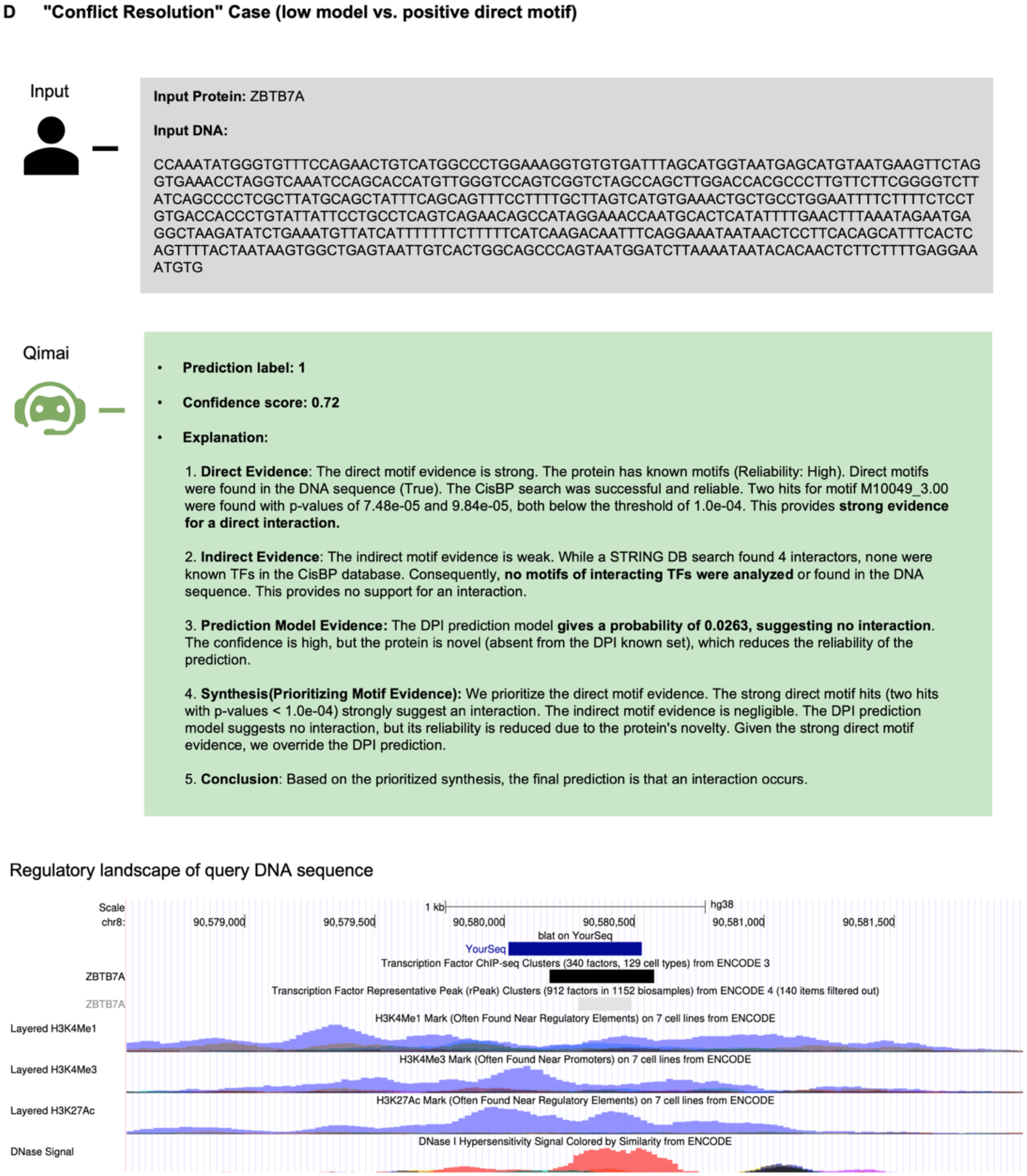
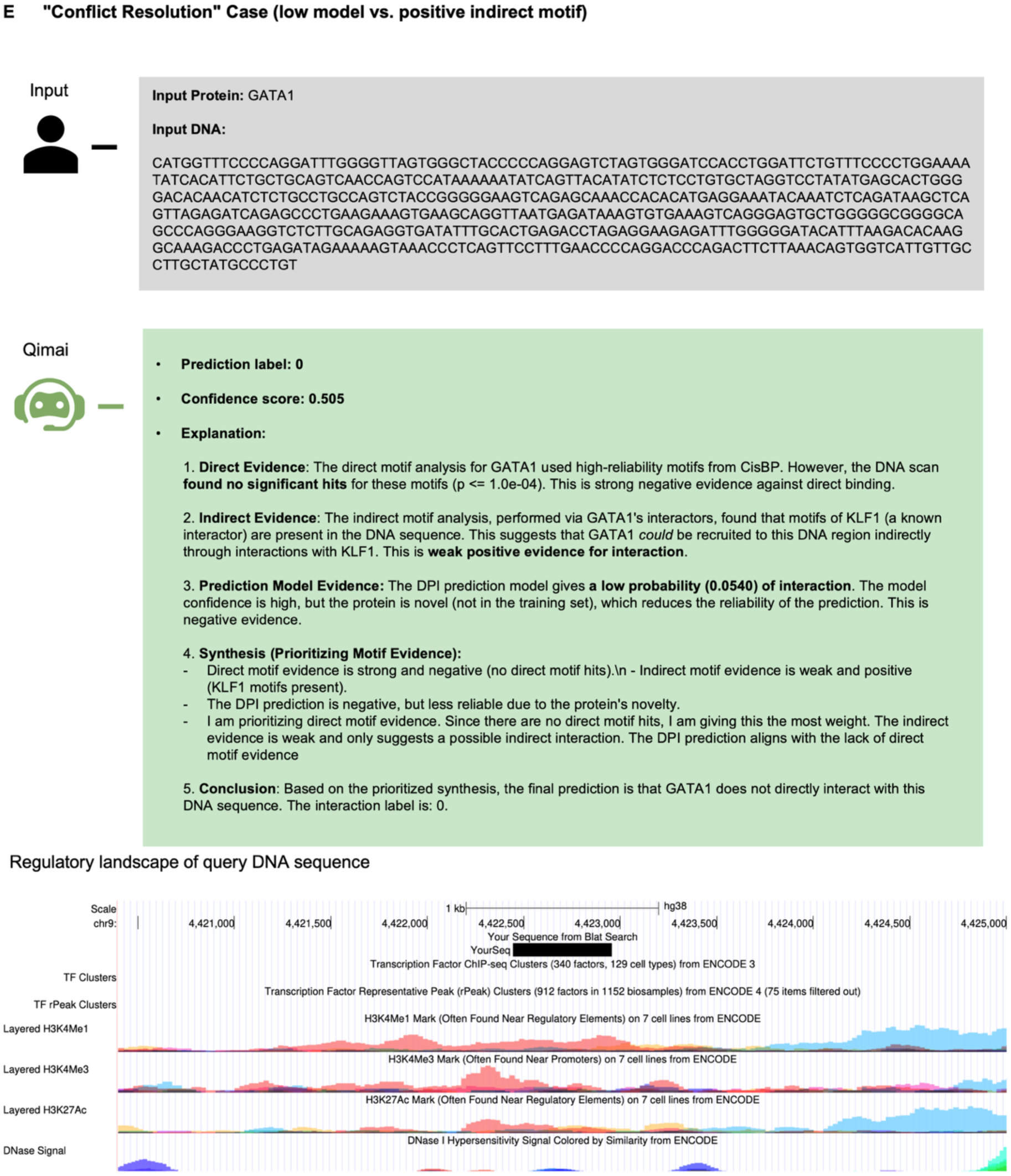
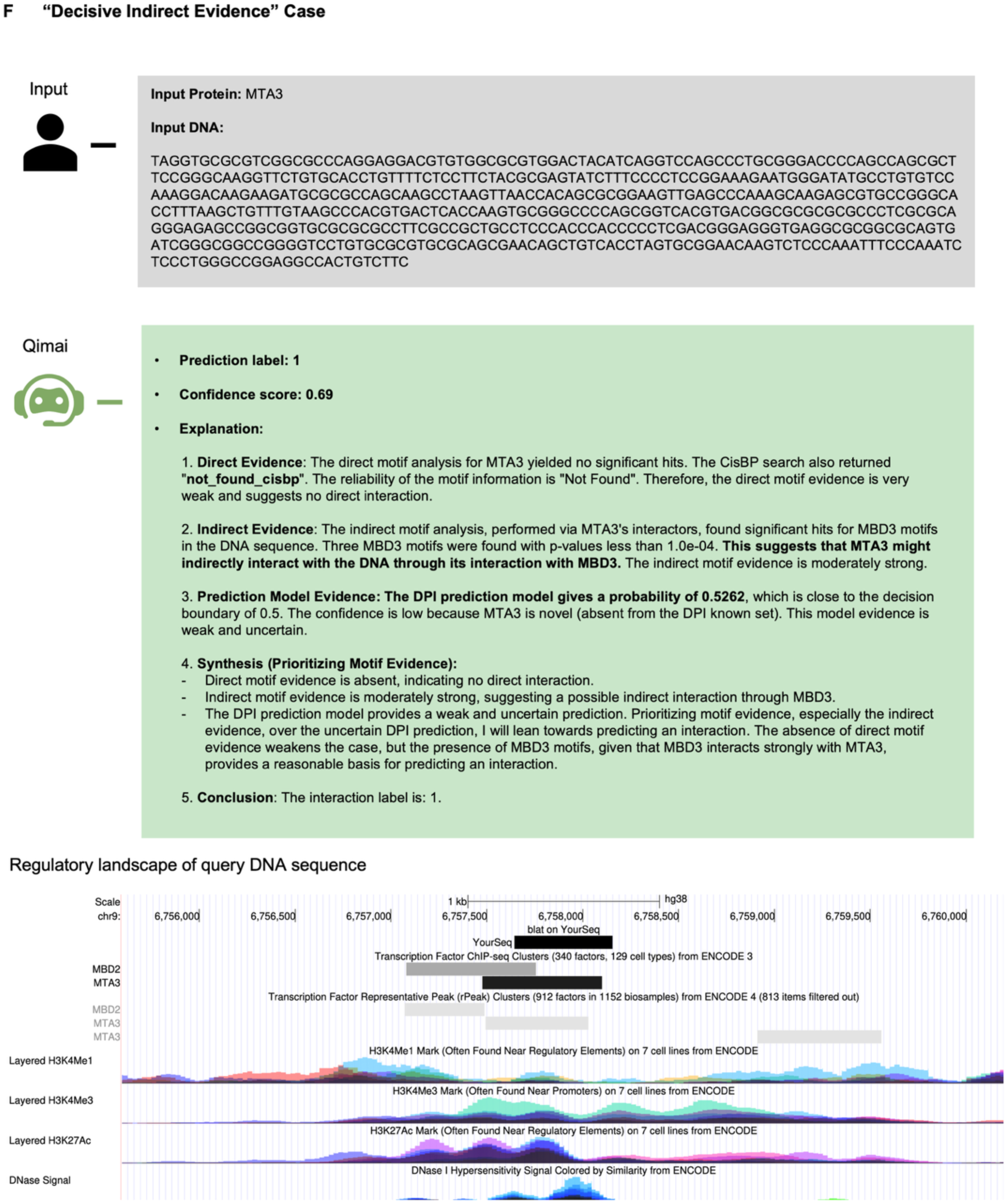
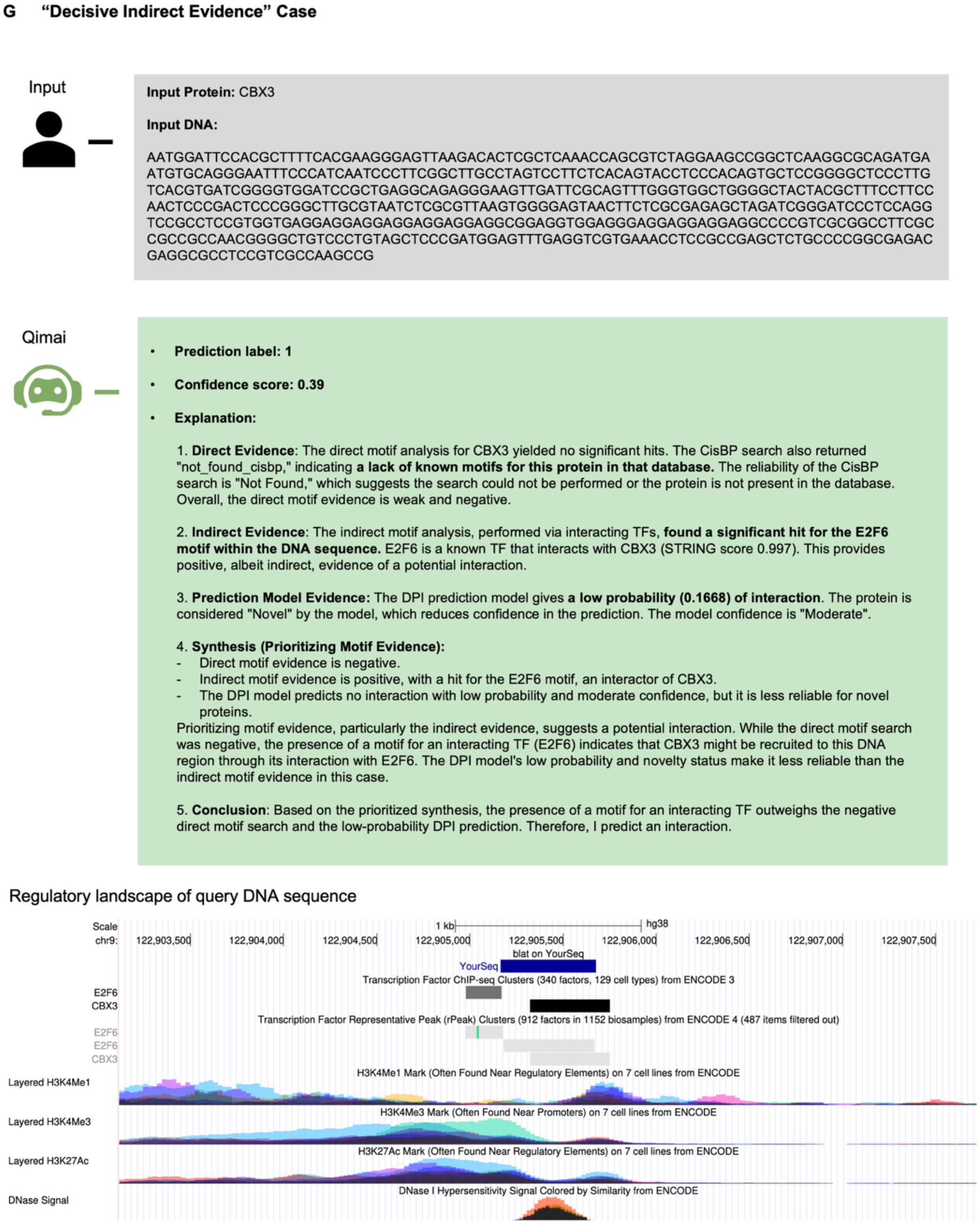
Representative case studies across diverse evidence scenarios. ChIP-seq binding peaks were used as ground truth. Genome browser view at the bottom shows the true ChIP-seq binding peaks along with histone marks and DNase signal. **(A) “All Signals Green” Case**. The TF SPI1 shows strong direct motif evidence, supportive indirect evidence, and a high DPI model score (0.7190), leading to a confident positive prediction. **(B) “Correct Rejection” Case.** The TF IRF4 lacks both direct and indirect motif evidence, which, combined with a low DPI model score (0.1890), results in a confident negative prediction. **(C) “Conflict Resolution” (low model vs. positive motifs).** For the TF BATF, strong direct and indirect motif evidence strongly suggests an interaction, prompting Qimai to override the conflicting low DPI model prediction (0.0255). **(D) “Conflict Resolution” (low model vs. positive direct motif).** The TF ZBTB7A only has direct motif evidence. Qimai prioritizes the presence of direct binding over the low DPI model score (0.0263) to predict an interaction. **(E) “Conflict Resolution” (low model vs. positive indirect motif).** For TF GATA1, weak indirect evidence from its partner KLF1 is insufficient to overcome the lack of direct motif evidence and a low DPI model score (0.0540), leading to a negative prediction. **(F) “Decisive Indirect Evidence” Case.** For MTA3, a component of a chromatin remodeling complex, there is no known motif documented in CisBP (thus direct evidence is absent) and the model score is uncertain (0.5262). However, moderately strong indirect evidence for its partner MBD3 leads Qimai to predict an interaction. **(G) “Decisive Indirect Evidence” Case.** The TF CBX3 has no known motif in CisBP and has a low model score (0.1668). Qimai correctly predicts an interaction by prioritizing the indirect evidence from E2F6, a known binding partner.

**Supplemental Figure 6:**
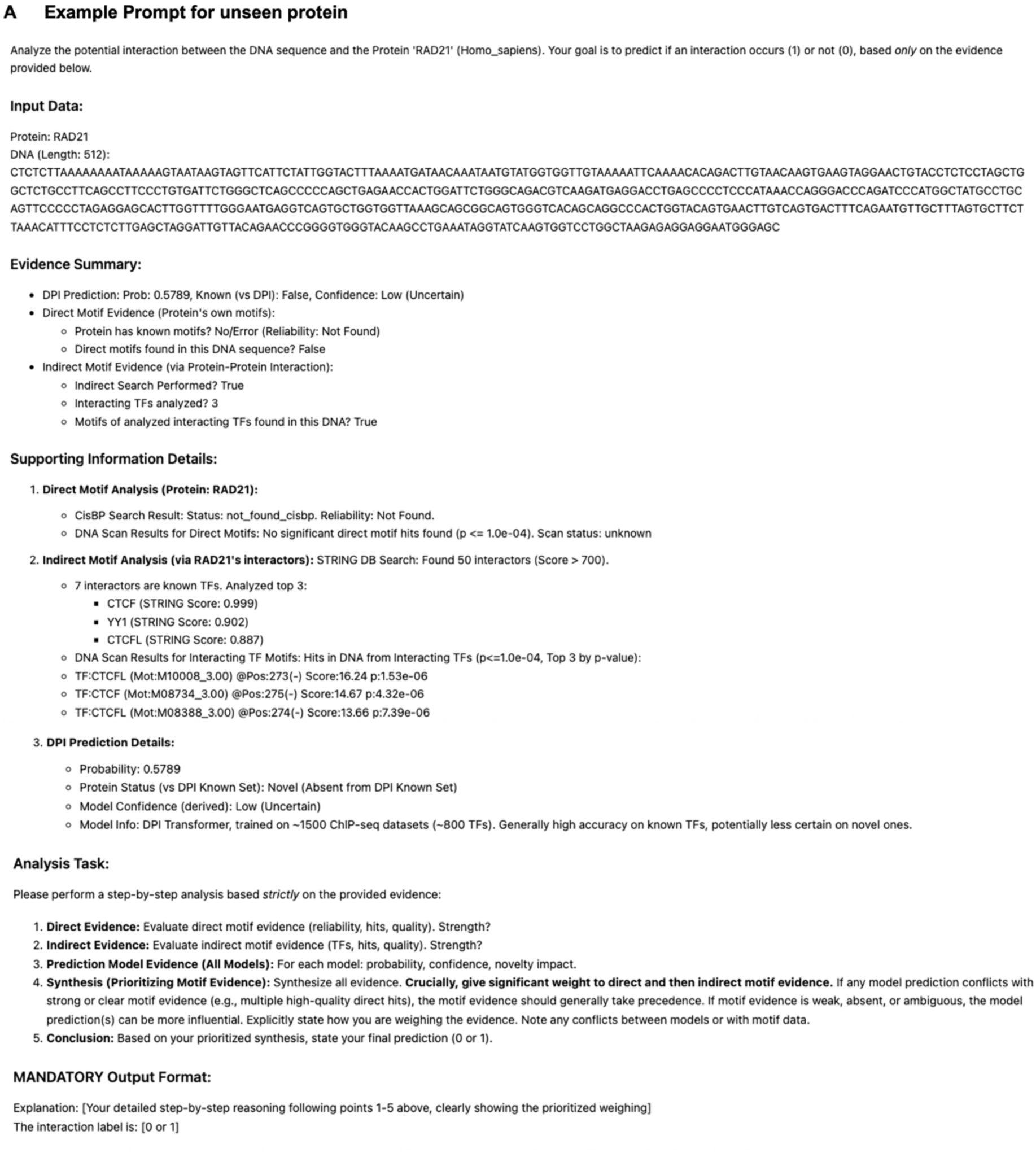
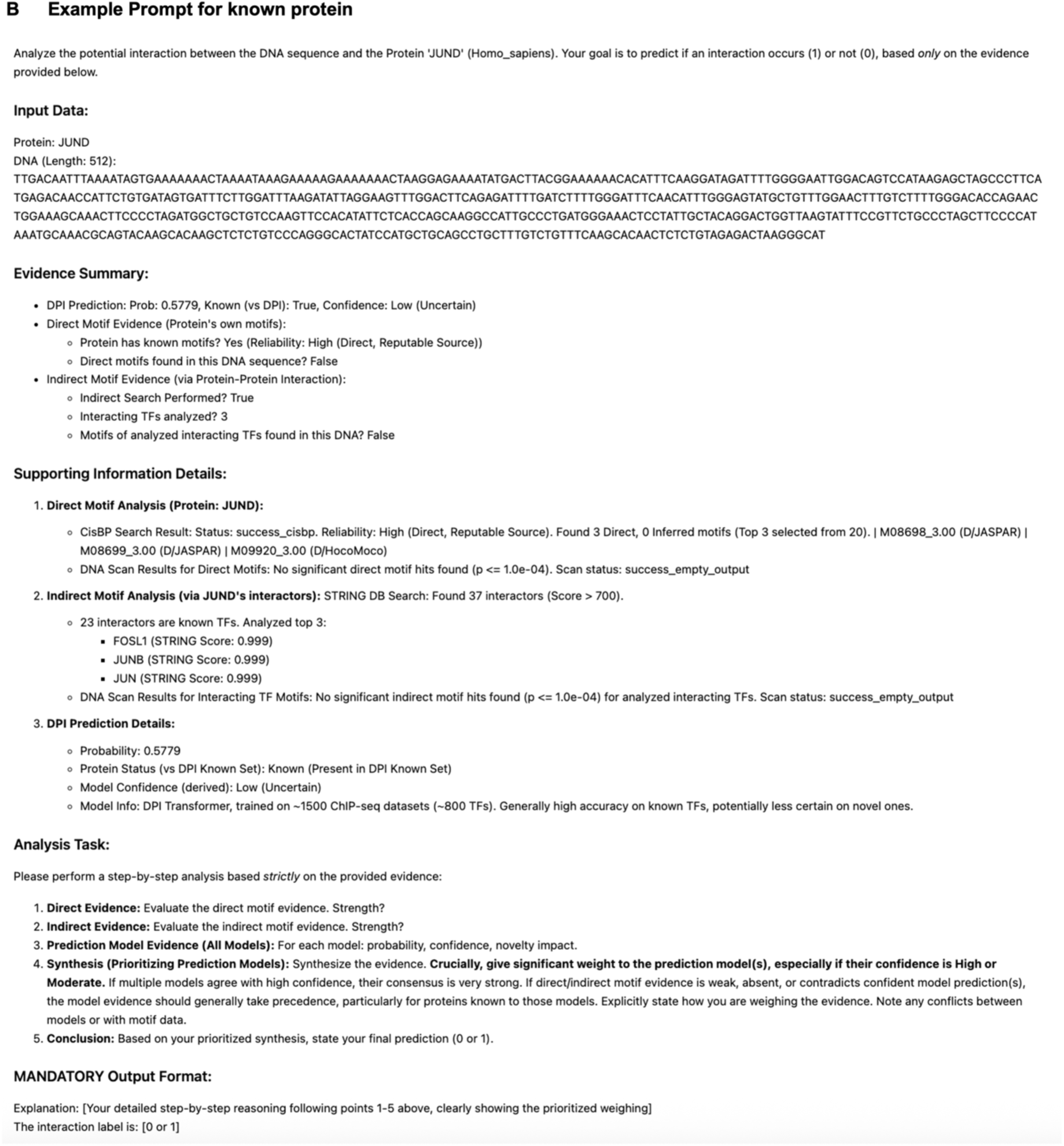
Prompts for reasoning LLM. **(A) Example prompt for unseen protein, which instructs LLM to prioritize biological evidence.** **(B) Example prompt for known protein, which instructs LLM to weigh the computational model prediction more.**

**Supplemental Figure 7:**
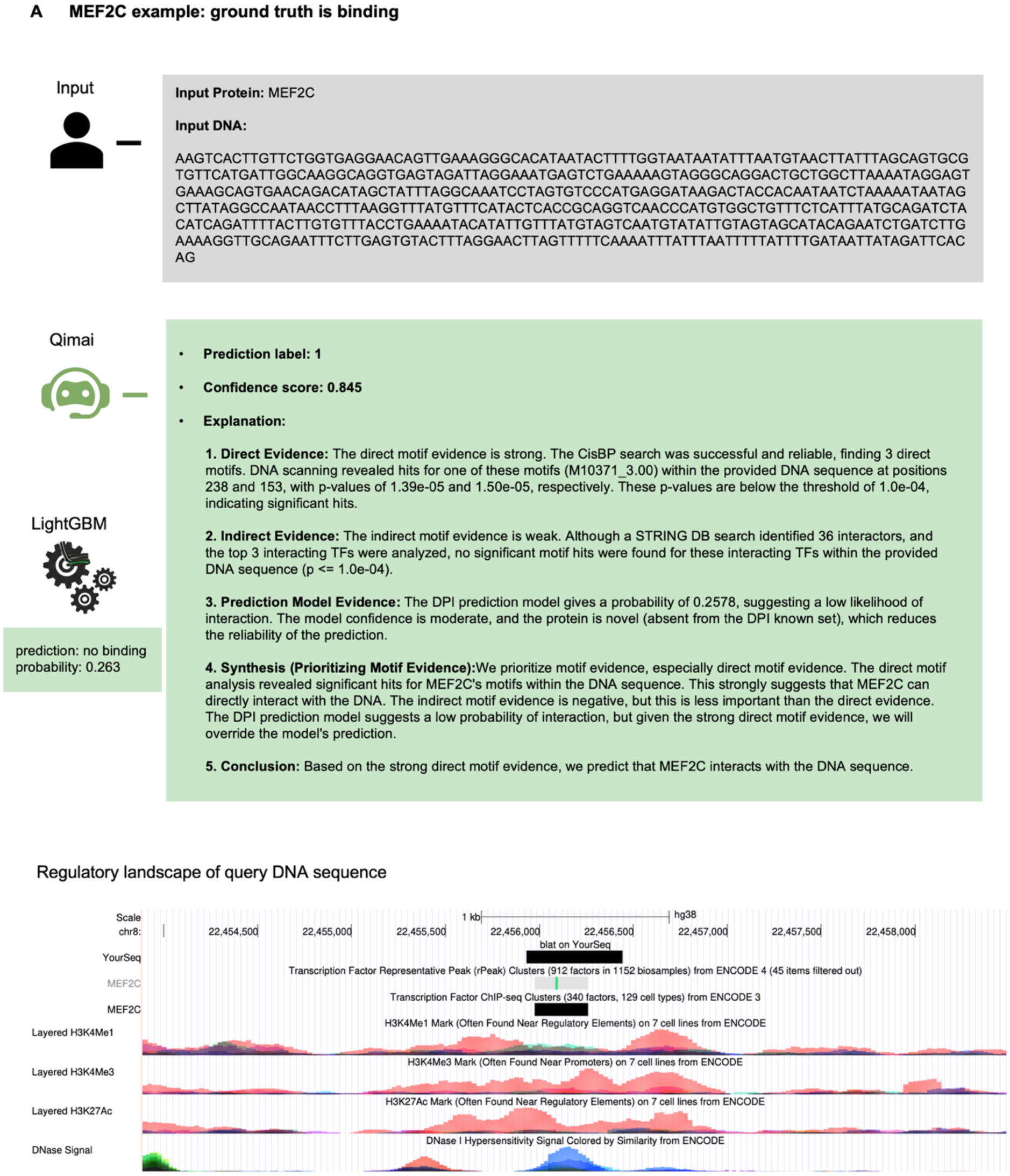
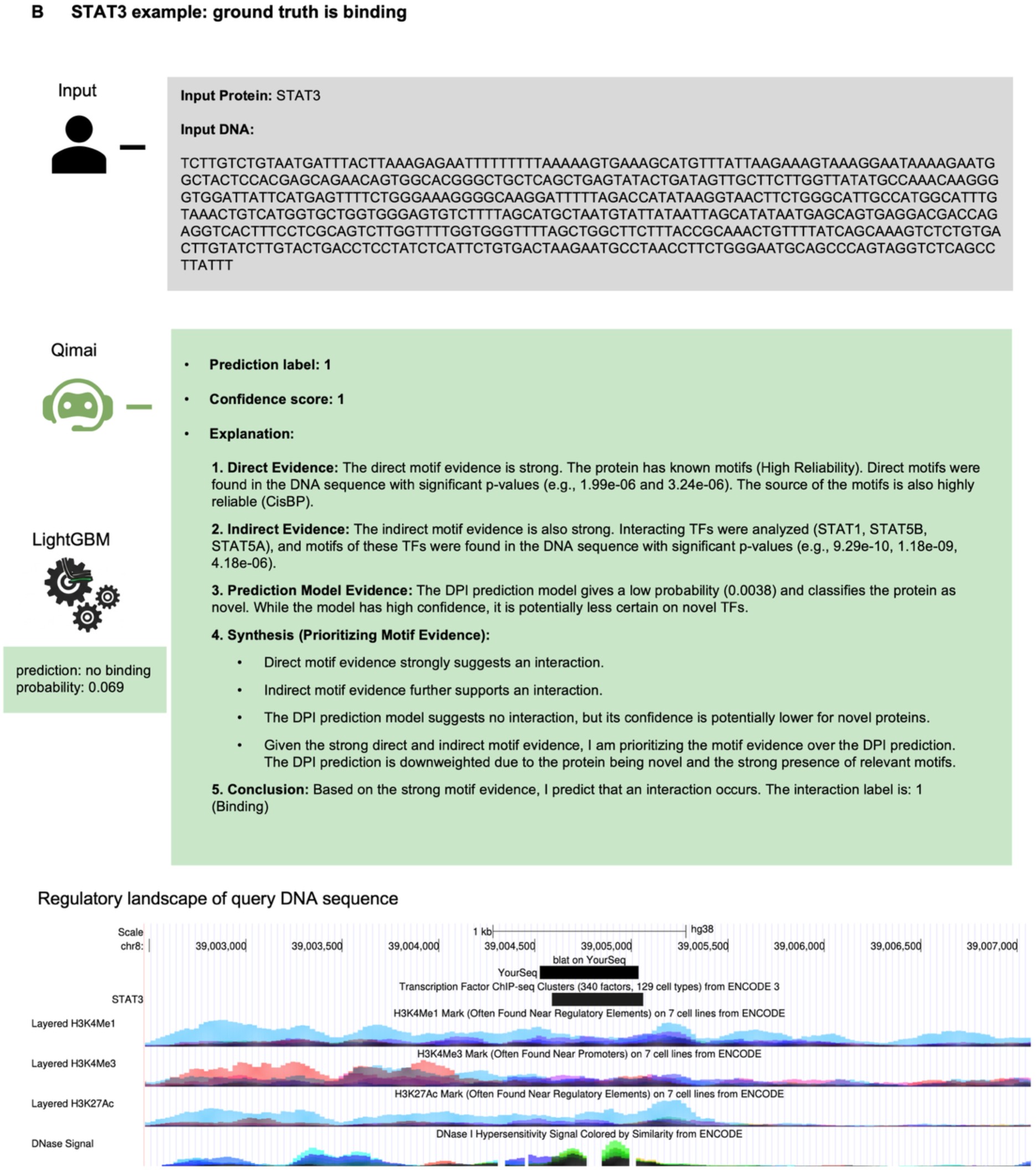
Representative examples in comparison with LightGBM classifier. **(A) MEF2C example.** It presents a clear conflict: strong direct motif evidence versus a low DPI model prediction (0.2578). The LightGBM classifier was misled by the low score and failed. Qimai, however, correctly prioritized the significant motif evidence over the unreliable model score for a novel protein, making the correct positive prediction, which is confirmed by ChIP-seq data. **(B) STAT3 example.** Despite overwhelming direct and indirect motif evidence, the DPI model returned an extremely low score (0.0338). The LightGBM classifier failed again by adhering to this score. In contrast, Qimai correctly identified the combined biological evidence as decisive, overriding the unreliable model to make the correct positive prediction, which is validated by prominent ChIP-seq peaks.

## Supplementary Notes

### A. Identifying and Mitigating DNA Sequence Bias in Dataset Preparation

Considering the similar performances of DPI-bench and DNABERT shown in **Figure 1C**, we investigated the contribution of protein module in DPI-bench. We removed the protein encoder and decoder blocks from the original model. Thus, the refined DNA embedding was directly fed into the prediction module to generate binary prediction. We trained the “DNA only” model on the same training set and evaluated the performance on the same testing set. Full model (DNA + protein) and DNA only model performed almost equally well (**Supplementary Figure 1D**), suggesting the protein module made negligible contribution to the model performance. DNA information plays an overwhelming role as compared to protein information, consistent with that DPI-bench model improved only slightly from DNABERT. Thus, DPI-bench model mainly learns how to classify different DNA sequences rather than different DNA-protein pairs, resulting that DNA patterns mislead the model to make the same prediction (binding or non-binding) for the same DNA sequence regardless of proteins.

To resolve the DNA sequence bias, we adopted a more rigorous data preparation process. In the previous dataset, the positive DNA sequences are genome sequences centered around ChIP-seq peaks identified from experiment while the negative DNA instances are synthetic sequences. Although the dinucleotide frequency is kept the same, the positive and negative sequences can have otherwise different patterns. Considering the main interest lies in the human genome in future inference, it is more appropriate to construct both positive and negative sequences from the genome.

Furthermore, each ChIP-seq experiment has its own unique set of peaks. It is unlikely to find exactly the same two peaks from different experiments even for the same TF. It can be problematic because each DNA sequence always has the same label regardless of the interacting protein, providing a lazy “shortcut” for the model, i.e. the model can make satisfactory predictions by just looking at the input DNA sequence. Thus, we need to construct instances with different labels associated with different proteins for the same DNA sequence. In this way, the model will be forced to utilize protein information to understand interaction modes and make opposite predictions for the same DNA. If a model only memorized the DNA patterns, the prediction accuracy would only be 50%, equal to random guess. Therefore, this label reversal design will help alleviate the DNA sequence bias.

Following DeepSEA protocol with a few adaptations (**Methods**), we re-processed the ChIP_690 datasets. The positive DNA sequences are genome segments overlapped with ChIP-seq peaks while the negative instances are genome segments without overlap of ChIP-seq peaks. Same DNA sequence can bind to some proteins (positive labels) and not bind to others (negative labels).

### B. Validation and Calibration of Qimai’s Confidence score

A critical component of Qimai is its ability to generate a final confidence score for its predictions. To be useful, this score must be not only discriminative (good at separating positive and negative cases, as shown by AUC-ROC/PR in the main text) but also well-calibrated, meaning the confidence level should correspond to the actual likelihood of a correct prediction. This supplementary analysis validates the calibration of Qimai confidence score against the raw probability output of a standalone DPI model, which serves as a primary input to our agent.

The results demonstrate a significant improvement of calibration by Qimai. As shown in the **Supplementary Figure S2B**, the standalone DPI model is markedly under-confident; for instance, in the lowest-confidence bin (predicted probability ≈ 0.1), the actual fraction of positives is more than 40%. Qimai corrects this behavior, producing a more confident score that tracks the ideal calibration line much more closely. The background histogram reveals that the majority of predictions fall into these lower-confidence bins, highlighting that the Qimai’s primary contribution is to improve reliability precisely where it is most needed. This validation confirms that the Qimai’s final confidence score is a trustworthy and interpretable measure of prediction accuracy.

To assess the robustness of the final confidence score, we evaluated the system’s performance on unseen data using four distinct sets of heuristic parameters (**Supplementary Table 11**). These configurations were designed to test different evidence-weighting philosophies: “Default” style prioritizing motif evidence, “V1” style that gives greater weight to computational model predictions, “V2” style with more conservative parameters (lower baseline confidence and smaller bonuses), and “V3” style with more optimistic parameters (higher baseline and larger bonuses). As shown in the **Supplementary Figure S3A (**left panel), key metrics remained highly stable across these different configurations. AUC-ROC was consistently high, ranging from 0.641 to 0.682, while AUC-PR was similarly stable, ranging from 0.679 to 0.701. This minimal variation demonstrates that the system’s overall predictive accuracy is not dependent on a specific, fine-tuned set of heuristic parameters.

Furthermore, we generated calibration plots to evaluate the reliability of the confidence score produced by each parameter set (**Supplementary Figure S3A**, right panel). The analysis shows that all four configurations yield well-calibrated probabilities, with their curves closely tracking the ideal diagonal line, which represents perfect calibration. This indicates that the confidence score is a reliable measure of the likelihood of a correct prediction, regardless of the underlying heuristic strategy. Taken together, the stability in performance metrics and the consistent calibration across diverse parameter sets provide strong evidence for the robustness of our confidence scoring architecture.

### C. Robust performance across different LLMs and different combination of prediction models

To further characterize the robustness of the Qimai framework, we performed additional analyses evaluating its consistency across different Large Language Models (LLMs) and its sensitivity to the combination of computational prediction models used as evidence.

A core component of the Qimai framework is its LLM-based Reasoning Agent. To ensure that the system’s strong performance is not dependent on a single proprietary model, we benchmarked Qimai’s performance using three distinct LLMs: Gemini-1.5-flash, Gemini-2.0-flash, and the open-source model Gemma-3-27b-it (**Supplementary Figure S3B**). The evaluation was conducted on both the seen proteins and the unseen proteins from external ChIP_690 dataset.

On the seen protein set, two Gemini-flash models enabled Qimai to achieve high performance while gemma-3 model showing a slight disadvantage. However, on the “unseen” dataset, which is a more critical assessment, all LLMs delivered comparable and strong performance, significantly outperforming standalone models. The open-source Gemma-3-27b-it model performed on par with the proprietary Gemini models across metrics like accuracy, F1-score, and AUC-ROC. This demonstrates that the Qimai framework’s architecture is robust and that its superior generalization capabilities are a product of its evidence synthesis design rather than an artifact of a specific LLM. This flexibility allows for the deployment of Qimai with powerful, locally-hosted open-source models, enhancing accessibility and reproducibility.

The Qimai framework is designed to synthesize evidence from one or more computational “Decision Agent” models. We conducted an ablation study to determine whether integrating predictions from multiple models simultaneously provides an advantage over using a single, best-performing model (**Supplementary Figure S3C**). We compared three system configurations: “DPI”, where only the DPI model’s prediction was provided to the LLM; “DPI + Sei”, where predictions from both DPI and Sei were provided; and “DeepSEA + DPI + Sei”, where predictions from all three models were provided. This comparison was performed on both the seen and unseen proteins.

On the seen protein set, integrating predictions from multiple models (“DeepSEA + DPI + Sei” and “DPI + Sei”) resulted in the highest performance across most metrics. This suggests that for in-distribution proteins, the LLM can effectively leverage the diverse predictive strengths of multiple models to achieve a more nuanced and accurate final prediction.

However, on the more challenging “unseen” dataset, this trend reverses. The configuration using only the single best-generalizing model (“DPI”) outperforms configurations that include predictions from Sei and DeepSEA across most of the metrics. This highlights a key insight: the optimal strategy for evidence integration depends on the novelty of the query. For out-of-distribution scenarios involving novel proteins, providing the single most reliable computational prediction minimizes potentially conflicting or less reliable signals from models that do not generalize as well, proving to be the most robust strategy.

### D. Representative examples of Qimai context-aware predictions

As stated in the main text, a key advantage of the Qimai is its flexible, context-aware reasoning, which allows it to outperform traditional classifiers when faced with conflicting evidence. While the main text highlights a case for the AP2A protein, the examples in **Supplementary Figure S7** for MEF2C and STAT3 further illustrate the robustness of the LLM Agent compared to a LightGBM classifier trained on the same evidence components.

The MEF2C case (**Supplementary Figure S7A**) presents a classic conflict between strong biological evidence and a misleading model prediction. The agent identified two highly significant direct motif hits for MEF2C, providing strong evidence for a direct binding event. However, the underlying DPI prediction model, being unfamiliar with this “novel” protein, produced a low probability score of 0.2578, suggesting no interaction. The LightGBM classifier, having learned a strong dependency on the model score during training, was misled by this low probability and incorrectly predicted no binding. In contrast, Qimai correctly reasoned that the model’s reliability is reduced for a novel protein and prioritized the high-confidence direct motif evidence, leading to the correct positive prediction. This conclusion is visually confirmed by the strong MEF2C ChIP-seq peak and active epigenetic marks in the provided genome browser view.

The example of STAT3 (**Supplementary Figure S7B**) offers an even more compelling demonstration. Here, overwhelming biological evidence was presented: multiple strong direct motif hits for STAT3, corroborated by strong indirect evidence from motifs of other STAT family members known to interact with it. Despite this, the DPI model returned an extremely low probability of 0.0338. The LightGBM classifier again failed decisively, adhering to the low model score and predicting no binding. Qimai, however, synthesized the full context. It identified the protein’s novelty, recognized the strength and consistency of both the direct and indirect motif evidence, and correctly concluded that this biological evidence far outweighed the unreliable model score. Therefore, it made a high-confidence positive prediction, which is validated by the ChIP-seq signals for STAT3 at that genomic location.

Together, these cases underscore the brittleness of fixed-weight classifiers in novel biological contexts and highlight the superior performance of the Qimai’s context-aware reasoning framework.

### E. Representative Case Studies of Qimai predictions

To provide a comprehensive view of the Qimai’s reasoning capabilities, we presented a few distinct DNA-protein interaction scenarios. These cases illustrate how Qimai synthesizes different types of evidence to arrive at a final prediction, ranging from straightforward confirmations to complex conflict resolutions.

**Supplementary Figure S5A** and **B** depict scenarios where all available evidence points in the same direction. In the “All Signals Green” case for the transcription factor SPI1 (**Supplementary Figure S5A**), Qimai’s positive prediction (Label: 1) is strongly supported by converging evidence. SPI1, also known as PU.1, is a well-established pioneer transcription factor that is essential for the development of myeloid and B-lymphoid cells. As a pioneer factor, it binds to specific DNA sequences called PU-boxes to open chromatin, making the presence of high-reliability direct motif hits a strong and expected indicator of interaction. This direct evidence, combined with supportive indirect evidence from known interacting partners like GATA1 and a high-probability score (0.7190) from the DPI model, makes for a high-confidence prediction. This conclusion is visually confirmed by the strong SPI1 and GATA1 ChIP-seq peak and active epigenetic marks in the provided genome browser view.

Conversely, **Supplementary Figure S5B** illustrates a “Correct Rejection” for the protein IRF4. IRF4 is another crucial transcription factor for the development and function of immune cells, including T-cells, B-cells, and macrophages. Its activity is highly context-dependent and often relies on cooperation with other transcription factors. In this specific instance, the lack of any significant direct motif hits for IRF4 or indirect hits for its partners aligns with the low interaction probability (0.1990) from the model. This leads to a confident negative prediction (Label: 0), correctly identifying this as a non-binding site. This conclusion is visually confirmed by the absence of IRF4 ChIP-seq peaks in the provided genome browser view.

A key strength of Qimai system is its ability to resolve conflicts between the initial computational prediction and biological evidence, particularly when a protein is novel to the prediction model. **Supplementary Figure S5C** and **D** showcase scenarios where the agent correctly prioritizes strong motif evidence over a conflicting low-probability score from DPI model. For instance, BATF (**Supplementary Figure S5C**), a member of the AP-1 family of transcription factors, is predicted to interact with the DNA despite a very low model score. BATF often functions by forming heterodimers with JUN family proteins to bind DNA. Although the DPI model returned a low probability (0.0255), likely due to the protein’s novelty to the model, Qimai identified strong direct motif evidence for BATF and, critically, indirect evidence for its interacting partners JUN and JUNB. By prioritizing the strong, biologically plausible direct and indirect motif evidence, Qimai correctly overrides the model’s initial prediction to conclude that an interaction occurs. This conclusion is visually confirmed by the strong BATF, JUN, JUNB, and IRF4 ChIP-seq peak and active epigenetic marks in the provided genome browser view.

**Supplementary Figure S5D** presents a complementary and even more stringent test with the protein ZBTB7A. Like BATF, ZBTB7A is a known transcription factor, but the DPI model, being unfamiliar with it, returned a similarly low probability of interaction (0.0263). The agent found two strong direct motif hits with highly significant p-values (7.48e-05 and 9.84e-05). Critically, in this case, there was no supporting indirect evidence. Despite the lack of secondary support, the agent’s logic remained robust. It identified the protein’s novelty as a reason to reduce the reliability of the model’s prediction and prioritized the strong, unambiguous direct motif evidence as the most trustworthy signal, once again overriding the model to correctly predict an interaction. This conclusion is visually confirmed by the presence of ZBTB7A ChIP-seq peaks in the genome browser view.

Qimai also demonstrates nuanced reasoning when direct evidence is absent. For instance, GATA1 (**Supplementary Figure S5E**) is a master regulator of red blood cell development. Here, Qimai was faced with conflicting signals: there were no direct GATA1 motif hits (strong negative signal), but weak positive evidence for a motif hit of its cofactor, KLF1, and the model score was low (0.0540). In this scenario, Qimai assigned more weight to the absence of direct hit and concluded that the weak indirect evidence was not sufficient to overcome this, resulting in a correct prediction of no interaction. This conclusion is visually confirmed by the absence of GATA1 ChIP-seq and active epigenetic peaks in the genome browser view.

**Supplementary Figure S5F** and **G** highlight scenarios where indirect evidence becomes the deciding factor, particularly for proteins that do not bind DNA directly. MTA3 (**Supplementary Figure S5F**) is a component of the NuRD chromatin remodeling complex, which does not bind DNA in a sequence-specific manner but is recruited to gene targets by other proteins. Here, direct motif evidence for MTA3 was absent, and the model’s prediction was uncertain (0.5262). However, Qimai found moderately strong indirect evidence of a motif for MBD3, another component of the NuRD complex. By prioritizing this strong biological clue of a known partner protein, it correctly predicts that an interaction occurs, likely through MTA3 being part of the recruited NuRD complex. This conclusion is visually confirmed by the MBD2 and MTA3 ChIP-seq peaks and active epigenetic marks in the genome browser view.

The case of CBX3 (**Supplementary Figure S5G**) further demonstrates this recruitment-based reasoning. CBX3 is a chromobox protein that reads histone modifications and lacks a known DNA-binding motif. As expected, the direct motif search failed, and DPI model produced a moderately low probability score (0.1668). However, Qimai identified a significant motif for the TF E2F6, a known high-confidence interaction of CBX3 and correctly reasoned that the presence of E2F6’s motif is strong evidence for recruitment. This conclusion is visually confirmed by the CBX3 and E2F6 ChIP-seq peaks and strong open chromatin signals in the genome browser view.

This provides a crucial contrast to GATA1 case ((**Supplementary Figure S5E**), where similar conflicting evidence led to an opposite prediction. The key difference lies in the biological role of the primary protein. For a sequence-specific transcription factor like GATA1, the absence of its own motif is strong evidence against an interaction. In contrast, for a protein like CBX3 that is known to be recruited, the lack of a direct motif is expected and does not constitute strong negative evidence. Therefore, the significant E2F6 motif becomes the decisive piece of evidence for CBX3, compelling a positive prediction, whereas the weaker KLF1 motif for GATA1 was insufficient to override the strong negative signal from its own missing motif.

